# Analyses of *Xenorhabdus griffiniae* genomes reveal two distinct sub-species that display intra-species variation due to prophages

**DOI:** 10.1101/2024.03.08.584182

**Authors:** Jennifer K. Heppert, Ryan Musumba Awori, Mengyi Cao, Grischa Chen, Jemma McLeish, Heidi Goodrich-Blair

**Affiliations:** Department of Microbiology, University of Tennessee at Knoxville, Knoxville, Tennessee, USA; Elakistos Biosciences, Nairobi, Kenya; Division of Biosphere Sciences Engineering, Carnegie Institute for Science, Pasadena, California, USA

**Keywords:** *Xenorhabdus griffiniae*, nematode-bacterium symbiosis, prophage, CRISPR loci, pangenome, bacterial subspeciation, insect toxins, entomopathogenic bacteria

## Abstract

Nematodes of the genus *Steinernema* and their *Xenorhabdus* bacterial symbionts are lethal entomopathogens that are useful in the biocontrol of insect pests, as sources of diverse natural products, and as research models for mutualism and parasitism. *Xenorhabdus* play a central role in all aspects of the *Steinernema* lifecycle, and a deeper understanding of their genomes therefore has the potential to spur advances in each of these applications. Here, we report a comparative genomics analysis of *Xenorhabdus griffiniae*, including the symbiont of *Steinernema hermaphroditum* nematodes, for which genetic and genomic tools are being developed. We sequenced and assembled circularized genomes for three *Xenorhabdus* strains: HGB2511, ID10 and TH1. We then determined their relationships to other *Xenorhabdus* and delineated their species via phylogenomic analyses, concluding that HGB2511 and ID10 are *Xenorhabdus griffiniae* while TH1 is a novel species. These additions to the existing *X. griffiniae* landscape further allowed for the identification of two subspecies within the clade. Consistent with other *Xenorhabdus*, the analysed *X. griffiniae* genomes each encode a wide array of antimicrobials and virulence-related proteins. Comparative genomic analyses, including the creation of a pangenome, revealed that a large amount of the intraspecies variation in *X. griffiniae* is contained within the mobilome and attributable to prophage loci. In addition, CRISPR arrays, secondary metabolite potential and toxin genes all varied among strains within the *X. griffiniae* species. Our findings suggest that phage-related genes drive the genomic diversity in closely related *Xenorhabdus* symbionts, and that these may underlie some of the traits most associated with the lifestyle and survival of entomopathogenic nematodes and their bacteria: virulence and competition. This study establishes a broad knowledge base for further exploration of not only the relationships between *X. griffiniae* species and their nematode hosts but also the molecular mechanisms that underlie their entomopathogenic lifestyle.

## Background

Buried in soils across the world is living white gold, a rich, but as yet under-utilized bioresource: *Steinernema* nematodes. These insect-killing roundworms have been found in 51 countries to date [1–6] and are profitable commercial products for the control of insect crop pests. In addition, they are colonized by microbes, including obligate symbiotic bacteria from the genus *Xenorhabdus*, that produce a battery of useful biomolecules [7]. To date, 31 *Xenorhabdus* species have been described [7–9] found in association with *Steinernema* nematodes, and the two species work in tandem to infect and kill insects and exploit the nutrient rich cadaver for the reproductive stage of their shared lifecycle.

Mechanistically, the nematode’s *Xenorhabdus* bacterium gut symbionts potentiate their insect-killing trait and serve as the primary food source for the nematode. In the non-feeding, host-seeking infective juvenile (IJ) stage of the nematode’s lifecycle, *Steinernema* nematodes house their *Xenorhabdus* symbionts in a specialized tissue of the anterior intestine known as the receptacle. After an IJ successfully enters an insect *via* natural openings such as spiracles, it defecates into the hemolymph, its *Xenorhabdus* bacteria, which by their secretion of insect toxins, immunosuppression and growth in the hemolymph, kill the insect [7]. The resultant insect cadaver is an enclosed nutrient-rich niche that both nematode and bacterium leverage to reproduce proliferatively. Nematode fecundity is enhanced by the consumption of *Xenorhabdus* [10], and *Xenorhabdus* defends the niche by secreting bacteriocins, antimicrobials and scavenger deterrents, which antagonize both microbial and invertebrate competitors [11–13]. Before their exit from the insect cadaver and entry into the surrounding soil, nascent IJ nematodes are specifically colonized by *Xenorhabdus* in the anterior intestine and ultimately, the receptacle [14].

Each *Steinernema* nematode is colonized in the receptacle by a specific *Xenorhabdus* species. However, some *Xenorhabdus* species associate with multiple *Steinernema* nematode hosts [7] suggesting a relatively fluid partnership landscape in which the molecular determinants of host-symbiont specificity across the two genera are still being defined. Key to this understanding is a robust comparative analysis of the phylogenetic relatedness and genomic content of *Xenorhabdus* isolates. Genome assemblies that are lowly contaminated, with high levels of completeness and greater than 50x coverage, are sufficient for bacterial species delineation [15], comparative genomics and identification and analyses of genes-of-interest. Indeed, insights into the *Xenorhabdus-Steinernema* symbiosis, bacterial speciation, and entomopathogenicity have been gained through such analyses of *Xenorhabdus* genomes. For example, comparing the degree of genetic similarity across whole genomes using digital DNA-DNA hybridization (dDDH) has led to the delineation of five novel *Xenorhabdus* species to date [8, 9, 16, 17]. Also, a comparative analysis of *Xenorhabdus bovienii* CS03 and SS-2004 genomes revealed that CS03 is more adapted to destroying microbial competitors than is SS-2004 but encodes fewer genes associated with entomopathogenicity [18]. Pangenome analyses of *X. bovienii* strains revealed intra-species content variation including of prophage origin, suggestive of strain adaptation to specific host environments [19]. The centrality of insect immune modulation to *Xenorhabdus* entomopathogenicity was highlighted by the fact that 21 of 29 analysed *Xenorhabdus* genomes are predicted to encode GameXPeptides [20] which are suppressors of insect immune pathways [21].

Of the >100 *Steinernema* nematode species described to date, only *Steinernema hermaphroditum* is a self-fertilizing hermaphrodite [22, 23]. This makes *S. hermaphroditum* particularly well-suited for rigorous nematode genetic studies, including those on the dynamics of transmission of *Xenorhabdus* bacteria from one generation of their host nematode to another [24, 25]. To lay the groundwork for such studies [26], we aimed to comprehensively analyse genomes of *Xenorhabdus griffiniae*, gut symbionts of *S. hermaphroditum* [23, 27]. We hypothesize that insights into the *X. griffiniae-S. hermaphroditum* symbiosis are attainable through detailed comparative genome analyses of *X. griffiniae* strains and their close phylogenetic relatives. Here, we report our studies in which we delineated two species of *Xenorhabdus* among the analysed strains, reconstructed their phylogenetic relationships with the rest of the *Xenorhabdus* genus, and analysed their pangenome and unique loci including prophages, CRISPR, and those encoding phage tail-like structures, secondary metabolites and insect toxins, the last of which was substantiated through insect mortality assays.

## Methods

### Bacterial genome sequencing

*X. griffiniae* HGB2511 and *Xenorhabdus* sp. TH1 (respective Genbank accession numbers for *16s rDNA*: MZ913116-MZ913125 and OR047834) were isolated from infective juvenile nematodes of *Steinernema hermaphroditum* CS34 and *Steinernema* sp. TH1, respectively, as previously described [16]. *X. griffiniae* ID10 was purchased from BacDive (DSM 17911). Cultures were inoculated in lysogeny broth (LB) stored in the dark (dark LB:0.5% yeast extract, 1% tryptone, 0.5% NaCl) and grown overnight under agitation at 30°C. Genomic DNA was prepared for sequencing by phenol extraction and spooling (*X. griffiniae* HGB2511), using the Qiagen DNeasy Kit per the manufacturer’s instructions with minor modification, (*X. griffiniae* ID10), or by both methods (*Xenorhabdus* sp. TH1). The Qiagen DNeasy Kit protocol was modified to prevent the viscous *Xenorhabdus* cell lysate from clogging the DNA-binding column by diluting the lysate with 10 mL of Qiagen QBT buffer before running it through the DNA-binding column by gravity. Purified DNA was sequenced using both short-read and long-read sequencing at the Millard and Muriel Jacobs Genetics and Genomics Laboratory, at the California Institute of Technology (for *X. griffiniae* HGB2511 and *Xenorhabdus* sp. TH1) or Novogene (for *X. griffiniae* ID10). For the *X. griffiniae* ID10 genome, short-read sequencing was performed using Illumina NovaSeq 150 base pairs (bp) paired-end short-read sequencing with library construction consisting of genomic DNA fragmentation, Illumina adapter ligation and PCR amplification, followed by size selection and purification, resulting in 33 gigabases (GB), of which 842 megabases (MB) were used for assembly. For primer-free long-read sequencing, library preparation consisted of size selection, adapter ligation and purification using Beckman Coulter AMPure XP beads. Sequencing was performed on an Oxford Nanopore (ONP) PromethION platform with base calling performed using Guppy software [28] with standard parameters. Prior to assembly, the ONP long-reads were filtered using Filtlong v0.2.1 resulting in 1.13 GB that was used in the assembly. Short- and long-reads were assembled using Unicycler v0.5.0 [29], resulting in a coverage of 432X. Sequencing of the TH1 and HGB2511 was done via a similar workflow. For each, 1.13GB and ∼0.74GB of bases were obtained from the Illumina and ONP runs, respectively. Likewise, a similar hybrid assembly method was also used, resulting in assemblies that were 406x and 496x for HGB2511 and TH1 respectively. Genome characteristics were determined via the PATRIC platform [30] (Table 1). Coverage was calculated by taking the total number of base pairs used in the assembly and dividing that by the genome size [31]. EvalG was used to determine the quality and completion assemblies [32]. Names and Genbank accession numbers of genomes used in this study are found in Additional file 2.

**Table 1.** Characteristics of genomes assembled in this study. Characteristics were identified using analysis tools on the Bacterial and Viral Bioinformatics Resource Center platform.

| Strain | Total length*<br>(bp) | Completeness<br>(contamination) | GC<br>content<br>(%) | Total CDS<br>(with functional<br>assignments) | tRNA<br>genes | rRNA<br>genes | Depth |
| --- | --- | --- | --- | --- | --- | --- | --- |
| ID10 | 4767870 | 100% (0.4%) | 43.91 | 4688 (70%) | 80 | 22 | 432x |
| HGB2511 | 4594889 | 100% (0.2%) | 43.78 | 4369 (72%) | 81 | 22 | 406x |
| TH1 | 3770596 | 99.6% (0.7%) | 43.48 | 3469 (78%) | 81 | 22 | 496x |

\* The total length was equal to the Contig N<sub>50</sub> (bp) since all three chromosome genome assemblies circularised.

### Tree generated using Bayesian inference

Phylogenetic analysis was performed as described previously [33]. Briefly, select *Xenorhabdus* (30) and *Photorhabdus* (1) species for which genomic sequences were publicly available were analysed using MicroScope MaGe’s Gene Phyloprofile tool [34] to identify homologous open reading frames (ORF) sets (homologs with at least 50% identity with synteny) which were conserved across all assayed genomes. Putative paralogs were excluded from the downstream analysis to ensure homolog relatedness, resulting in 1235 homologous sets (one-to-one orthologs). Homolog sets were retrieved via locus tag indexing using Python v3.8.0, individually aligned using Muscle v3.8.31 [35], concatenated using Sequence Matrix v1.9 [36], and trimmed of nucleotide gaps using TrimAL v1.4 [37]. A General Time Reversible + γ variation substitution model was used for maximum likelihood and Bayesian analysis. Maximum likelihood analyses were performed via RAxML v8.2.10 [38] using rapid bootstrapping and 1,000 replicates and were visualized via Dendroscope v3.6.2 [39]. Nodes with less than 40% bootstrap support were collapsed. Bayesian analyses were performed via MrBayes v3.2.7a in BEAGLE [40] on the CIPRES Science Gateway platform [41]. A total of 100,000 Markov chain Monte Carlo (MCMC) replicates were performed. Twenty-five per cent were discarded as burn-in, and posterior probabilities were sampled for every 500 replicates. Two runs were performed with three heated chains and one cold chain. The final average standard deviation of split frequencies was 0.052352. Bayesian trees were visualized with FigTree v1.4.4 [42]. Posterior probabilities are 100% except where otherwise indicated.

### Digital DNA-DNA hybridisation and pangenome analyses

To determine pairwise digital DNA-DNA hybridisation (dDDH) values among 31 strains (ID10, VH1, XN45, TH1, HGB2511, BMMCB, BG5, Kalro, 97 and 22/26 validly published *Xenorhabdus* type strains), their fasta formatted genomes were uploaded to the TYGS server [43] and analysed as previously described [16]. Genomes of six *X. griffiniae* strains plus that of *Xenorhabdus* sp. TH1 were comparatively analysed using a pangenome approach in Anvio 7.1 [44]. Briefly, fasta formats of the seven genomes were reformatted to simplify the definition lines, then converted to anvio contig databases. On these, hidden Markov models (HMMs) and genes were identified using HMMER [45] and Prodigal [46], respectively. The functions of these genes were then predicted, based on orthology, using the Cluster of Orthologous Genes (COG) database [47] as a reference. These annotated contig databases were then used to construct a pangenome with the anvi-pan-genome program under the following parameters --use NCBI-BLAST, MCL inflation 10, minbit 0.5, --exclude-partial-gene-calls. The anvi-display-pan program was used to both display the pangenome as a sunburst chart and subsequently create selected bins. To obtain sub-pangenomes of only type VI secretion system-associated orthologous gene clusters (GC), all GCs annotated with “type VI” were identified using the search functions feature and binned. The anvi-split program was then used to obtain a pangenome of only Type VI secretion system-associated GCs (Supplementary Sheet S3 in Additional File 2). For each analysis, pangenomes were represented both as sunburst charts and tabulated gene lists. To calculate average nucleotide identities (Supplementary Sheet S13 in Additional file 2), the fastANI [48] program was used within an Anvio environment. The average alignment fraction and fragments were 0.85 and 1425 respectively.

To identify the mobilome, all genes annotated with the COG category “X” were extracted from the main pangenome and used to create a sub-pangenome of the mobilome only. This was then used to calculate the total number of genes annotated as phage, transposase, and plasmid-related in each genome. For each genome, these totals were correlated with proteome size using Pearson’s adjusted (due to the small sample size) r-square at a 0.05 alpha level.

### Identification and analysis of unique genes within the accessory genome

Using the tabulated output of the Anvio pangenome analysis and Microsoft Excel, core genes and genes unique to a given strain were identified using the number of genomes in which the gene cluster has hits identifier, where the value equalled 7 for core genes (19,196) and 1 for unique genes for the HGB2511 (340), ID10 (411) and TH1 (454) strains. Because the genomes of the sub-species clade which includes *X. griffiniae* Kalro are so similar, there were less than ten unique genes per strain [XN45(8), VH1(7), xg97(4), Kalro (0)]. Thus, to gain a better understanding of what genes might be unique within the sub-species clade, we compared the Kalro genome alone to the HGB2511, ID10 and TH1 genomes and found 454 unique genes for further analysis. To elucidate the potential functional categories of genes enriched among those unique to a given species, the COG annotations assigned as part of the pangenome analysis were used. The number of unique genes per COG category was plotted as a percentage of the total number of unique genes with a COG designation for each strain and the core genome using GraphPad PRISM v10.1.1. As expected, a much larger fraction of genes with no COG designation was found in each of the unique gene categories (approximately 50% in each case) compared to the core genome (6%).

### Elucidation and analysis of prophages in *X. griffiniae* genomes

To identify prophages in bacterium genomes, we used VIBRANT 1.2.1 [49] under default parameters. Each prophage sequence was then separately reannotated with Pharokka [50] under “meta mode” and Bakta [51]. Genomad [52] was used to both taxonomically classify prophages and assess their quality (through CheckV [53]) and completion (Supplementary Sheet S14 in Additional File 2). Resultant gene lists of the annotated prophages are available in (Supplementary Sheets S8-S11 in Additional File 2).

To identify similar prophages across strains, we used progressive Mauve [54] to identify collinearity blocks between prophage sequences. Considerably similar pairs were selected, pairwise aligned, and visualised as dot plots using Geneious 8.1.9 [55].

To determine the effect of prophages on the dDDH values among *X. griffiniae* strains, all prophage sequences were deleted from their corresponding bacterium genomes in Geneious 8.1.9. Then, dDDH analyses were rerun using both original and “phageless” *X. griffiniae* bacterium genomes (Supplementary Sheet S12 in Additional File 2).

To identify strain-specific and subspecies-specific genes that are from prophages, a pangenome approach was used. Briefly, their entire prophage sequence from a strain — ID10, Kalro, TH1, xg97 and HGB2511— were merged into a fasta file. For example, all prophages from the ID10 genome were merged into a single fasta file “ID10_prophages.fasta”. These five prophage fasta files, plus those of genomes of ID10, HGB2511, VH1, XN45, xg97 and Kalro, were used to create a pangenome as aforementioned. Using the anvio-display-pan program, strain-specific or subspecies-specific GCs that were of prophage origin were visually identified and binned. Using the anvio-summary program, the number, annotation, and aa sequences of these strain-specific or subspecies-specific genes of prophage origin were obtained from resultant gene lists (Supplementary Sheet S14 in Additional File 2). This was then used to calculate what proportion of strain-specific and subspecies-specific genes were from prophages. To identify genomic loci encoding complete type VI secretion systems (T6SS) within genomes, we used Secret6 [56] under default parameters. Geneious was used to create multiple sequence alignments of T6SS-encoding loci and to calculate pairwise nucleotide sequence identities.

To infer gene gain and loss events within the *X. griffiniae* clade, the resultant pangenome of the seven strains was used to manually construct a phyletic pattern (Supplementary Sheet S6 in Additional File 2). A dDDH-based phylogenomic tree of six strains of *X. griffiniae* plus *Xenorhabdus* sp. TH1 only was reconstructed via the TYGS pipeline. Both this tree and the phyletic pattern were used as input data in COUNT [57]. The evolution of gene content was inferred using Wagner parsimony with a gain penalty of two.

### Identification and analysis of CRISPR-Cas systems in *X. griffiniae* genomes

To identify CRISPR-Cas systems in *X. griffiniae* genomes we used BlastN on the Magnifying Genome platform to search for CRISPR repeats with the previously identified *X. nematophila* CRISPR repeats, XnCRISPR-E and XnCRISPR-G as queries [58]. Regions with similar or identical sequences were extracted and manually curated for repeat-spacer content. All results were then verified using CRISPRDetect [59]. The only distinction between the two approaches was for ID10, for which the manual approach had suggested five potential repeat sites of one spacer only, whereas CRISPRDetect did not identify any. We therefore conducted a second manual annotation of the repeats in this genome (Additional File 3). Based on conservation with consensus repeats called by CRISPRDetect in both upstream and downstream repeats, we considered ID10 1ai and ID10 1bii regions to be bona fide single spacer CRISPR loci. Cas protein-encoding genes were identified based on their annotation in the Magnifying Genomes platform and confirmed using CasFinder-3.1.0 [60]. To identify protospacers we searched for full-length identical sequences using each identified spacer as a BlastN query against the *X. griffiniae*, TH1, and BMMCB genomes in the Magnifying Genome platform. Protospacers were identified in both other strains and within the same genome (self-targeting protospacers). To gain a broader view of the non-self protospacers, we used CRISPRTarget (http://crispr.otago.ac.nz/CRISPRTarget/crispr_analysis.html) to search the spacer sequences found by CRISPRDetect for TH1, ID10, HGB2511, and BMMCB genomes, against subset of the available databases (ALCAME genes, Genbank-Phage, RefSeq-Archaea, RefSeq-Plasmid, and RefSeq-Viral) for high confidence sequence matches to the potential protospacers [59, 61]. All putative protospacers for a given spacer were listed (Table S5 in Additional File 3) and the top, annotated hit for each is listed in Table 2.

**Table 2:**
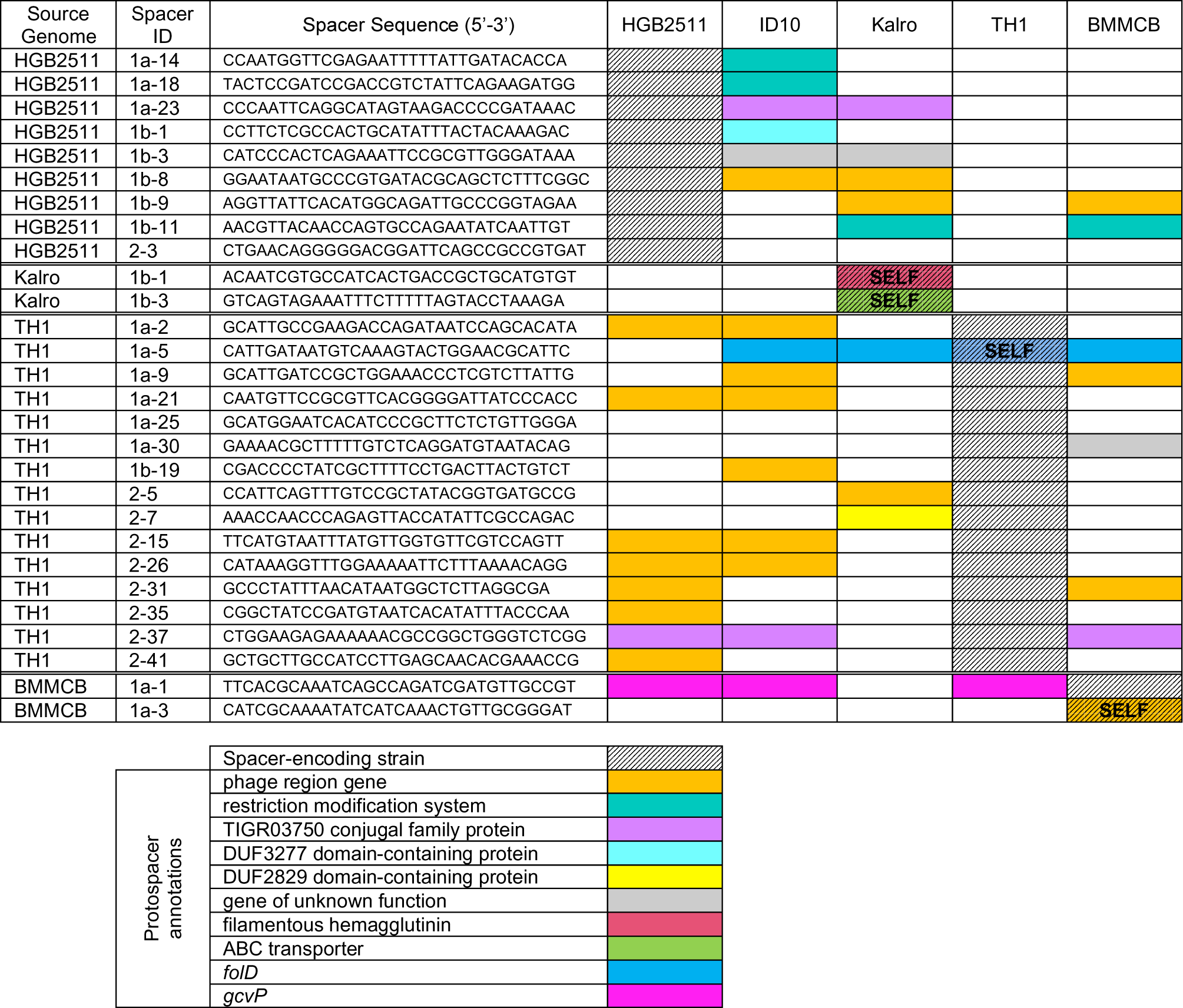
Annotated spacer-protospacer content among *X. griffiniae* and related strains.

### Elucidation of biosynthetic gene clusters

To predict which secondary metabolites can be produced by *X. griffiniae,* a fasta format of the ID10 genome was uploaded to the AntiSMASH [62] webserver (https://antismash.secondarymetabolites.org) as data input. Default parameters (i.e., relaxed detection strictness) and the use of all extra analyses were selected. Output data comprised detailed bioinformatic analyses of 23 biosynthetic gene clusters (BGCs) in the ID10 genome. Each of the 23 BGCs was then manually inspected in two main ways. First, the clusterBLAST feature was used to identify known homologous BGCs by comparing the gene synteny and sequence similarity of an ID10 BGC to those of known BGCs found in the MiBIG database [63]. An ID10 BGC was considered homologous to a known BGC if it contained >80% genes in perfect synteny, with each gene having BLASTp sequence similarity >47%, sequence coverage >40%, and E_value_ <2.78E-19 with the corresponding gene in the known BGC. Second, for ID10 non-ribosomal peptide synthetase (NRPS) BGCs, their Stachelhaus codes [64] for adenylation domains, as well as their epimerisation/dual condensation domains [65] were analysed to determine the amino acid sequence of the linearised non-ribosomal peptide (NRP) the NRPS was predicted to biosynthesize. Predicted NRPs were then compared to known NRP to identify NRPS BGCs that encode the biosynthesis of novel derivatives/peptides. This workflow was similarly applied for the analysis of BGCs in the HGB2511 genome. Chemical structures of the compounds whose production was predicted to be encoded by the known ID10 BGCs were obtained from the Natural Product Atlas [66] and edited in Chemdraw and Inkscape [67].

### Putative toxin, secretion system, Cas protein, and Restriction modification system identification

Putative toxins predicted to impact insect or nematode virulence were identified using multiple approaches. The first approach used the loci of previously characterized known or suspected toxins [68]. BlastP was performed on the Magnifying Genomes platform, and the BLAST query accession proteins are listed in the putative toxins table (Table 3), this procedure was repeated for other novel putative toxins identified as well. The PathoFact software package [69] was used to identify novel putative toxins in the ID10, HGB2511, Kalro, TH1 and *X. nematophila* 19061 genomes with the standard settings and chromosomal genomes downloaded from the Magnifying Genomes platform. The toxin library outputs from PathoFact were compared and toxins potentially unique to each genome were further examined (Additional File 4). This resulted in the identification of a potential hydrogen cyanide synthetase locus (XTH1_v2_1430-1432) which appears to be unique to the TH1 genome among those analysed. Further, this search revealed two zonula occludens toxin proteins (JASDYB01_14222 and JASDYB01_14237 and _14239) which appear to be unique to the Kalro genome (homologs of were also found in VH1, XN45, and xg97). To both confirm and expand the list resulting from the combination of the above analyses, the search term “toxin” was used on the Magnifying Genomes platform to further query the gene annotations for the HGB2511, ID10, TH1 and Kalro genomes. This search resulted in the identification of a protein annotated as an insecticidal toxin (XGHID_v1_0629) in the ID10 genome and a homolog (XGHIN1_v1_3228) was subsequently found in the HGB2511 genome via BlastP. The products of toxin-antitoxin systems were generally not considered in our analysis. The highest confidence and most interesting results are summarized in the Putative Toxins Table (Table 3).

**Table 3:**
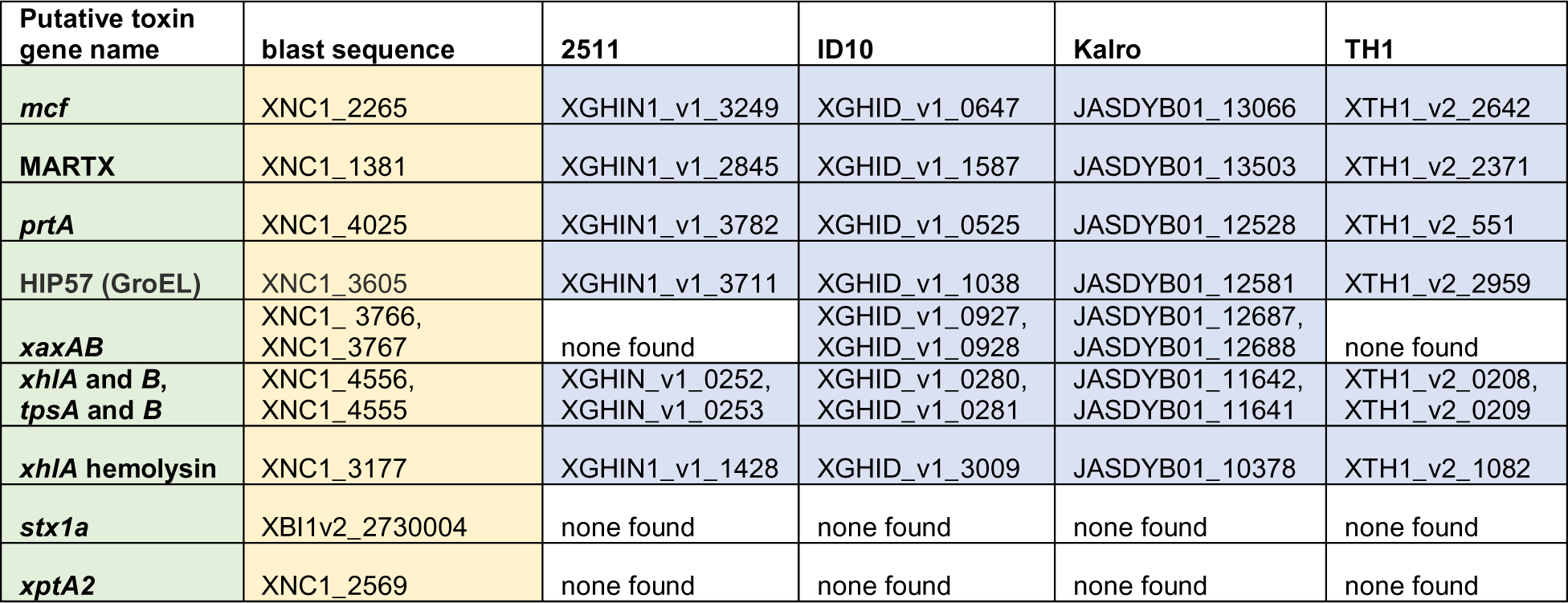
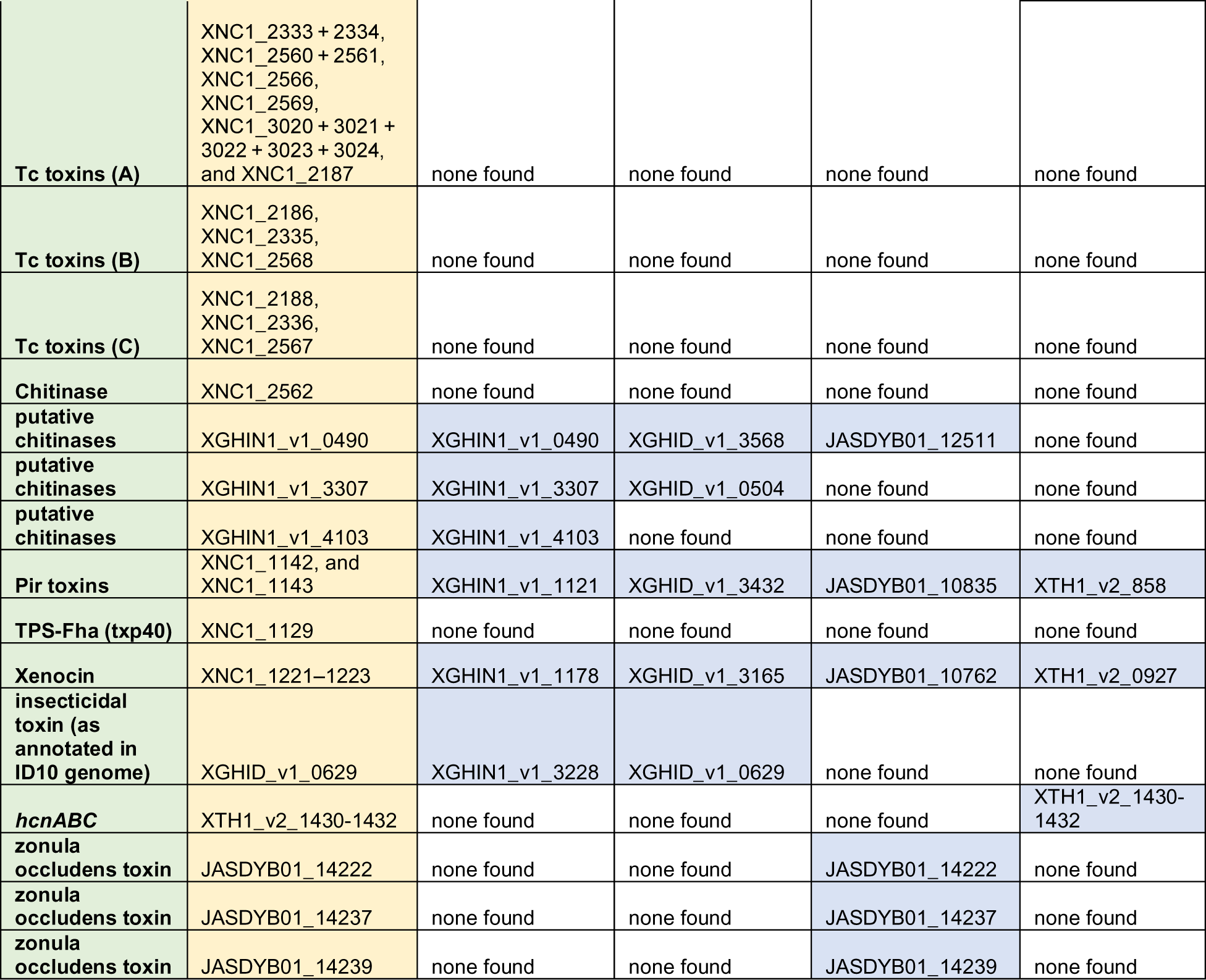
Putative entomotoxin-encoding genes in *Xenorhabdus griffiniae* and TH1 genomes.

To systematically predict the types and numbers of secretion systems encoded by the *X. griffiniae* genomes (HGB2511, ID10, Kalro, VH1, XN45, xg97), TXSScan application from the Macsyfinder 2.0 [70] program was run on the Galaxy [71] platform specifying an ordered, circular replicon, diderm bacteria and with default HMMER options. A combination of the summary output for each genome is found in Table S1 in Additional File 1. Alignment of MARTX regions was performed using MUSCLE in MegAlignPro (DNA Star).

To confirm the type and numbers of Cas proteins encoded by the *X. griffiniae* genomes (HGB2511, ID10, Kalro, VH1, XN45, xg97), CasFinder application from the Macsyfinder 2.0 [70] program was run on the Galaxy Pasteur platform specifying an ordered, circular replicon, diderm bacteria and with default HMMER options. The summary table is compiled from the software summary output and the additional sheets contain the best solution predictions from the Cas Finder output and include the gene names and locus tags from the different Cas systems found (Supplementary Sheet S20 in Additional File 2).

Restriction modification systems were identified by searching the BMMCB, TH1, ID10, HGB2551 and Kalro genomes for restriction enzymes, anti-restriction proteins, and restriction enzyme-associated methylases-based gene annotations on the MAGE Microscope platform. These genomes were further searched using BLAST 2.15.0+ to compare a list of “Gold Standard” methyltransferase and endonuclease protein sequences from the New England Biolabs’ REBASE with the protein sequences contained within each genome [72]. An E value cut-off of ≤1 × 10^−5^ and 75% coverage were used to generate a list of high confidence candidates. This candidate list was compared with the initial list of restriction system proteins generated using MAGE, and redundant sequences were removed. Methyltransferase homologs that were not found to be near predicted restriction endonucleases were excluded from further analysis, though the presence of many ‘orphan’ methyltransferases may indicate a need to protect the bacterial chromosome from restriction modification systems [73]. Putative restriction enzyme loci were examined to identify or confirm neighbouring methylases in the case of Type I, II and III restriction enzymes, or lack thereof, in the case of the Type IV and the HNH restriction endonucleases identified [74]. Of the putative restriction modification genes examined, only three loci from the BMMCB genome (LDNM01_v1_10020, LDNM01_v1_400040, LDNM01_v1_1980001) were predicted to encode restriction endonucleases from Type I or III but were not observed to encode a proximal methyltransferase. Because of their incomplete nature, these loci were excluded from the final table and count of restriction systems was identified. However, each locus was near a contig break, so a less fragmented BMMCB genome assembly may reveal these to be complete predicted restriction endonuclease loci.

### Insect larvae rearing and preparation

Eggs of *Manduca sexta*, the tobacco hornworm, were purchased from Carolina Biological Supply (North Carolina, USA) and were reared to the fifth instar according to a previously described protocol [75]. Briefly, eggs were sterilized with 0.6 % (v/v) bleach solution on arrival and then transferred to a four-ounce plastic container with a Gypsy moth diet (MP Biomedicals, Ohio, USA). The eggs were then incubated at 26 °C in a humidified insect incubator with a 16-h light: 8-h dark photoperiod. Once hatching was complete, the larvae were transferred to a new four-ounce container for two days and then transferred to a new two-ounce container. Feeding and cleaning were performed every two to three days until the larvae reached the fifth instar stage. On the day of the experiment, the larvae were examined and sorted by weight ranging from 0.67 g to 3.5 g. Larvae (n=10, with n=5 for phosphate-buffered saline (PBS) control) were placed into individual two-ounce plastic containers. Groups of 10 larvae were carefully injected with 10 µL of various doses (10^-5^, 10^-4^, and 10^-3^) of bacteria between the first set of abdominal prolegs using a Hamilton syringe. Following injections, the larvae were incubated at 26 °C in a humidified insect incubator with a 16-h light: 8-h dark photoperiod and monitored for survival over 72 h.

### Preparation of bacteria for infection of *M. sexta*

Bacterial strains were streaked from –80 °C freezer stocks onto dark LB agar and cultivated at room temperature in the dark for two days. Broad streaks including multiple colonies were used to inoculate LB medium (5 mL) and incubated on a rotating wheel at 30°C for approximately 8 h. At 8 h the optical density (OD) at 600nm was measured (OD600), and cultures were all near an OD600 of 1.0. Cultures were normalized to an OD600 of 1.0 by adjusting the volume of culture taken (e.g., 500 µL OD600 of 1.0). Cells were spun down for 1 min at max speed in Eppendorf tubes and washed twice with 1 mL of sterile PBS. After the final wash, cells were resuspended in 500 µL of PBS. The washed cells were then diluted ten-fold, six times in sterile PBS in a 96-well plate. For each dilution, 10 µL was inoculated onto LB agar plates (LBP) to quantify the number of colony-forming units (CFU) in each dilution. To test for sterility, PBS was also inoculated onto LBP. Based on a previous study [68], we estimated that 100, 1,000, and 10,000 CFU/10 µL was suitable for an observable virulence (insect mortality) and dose-response, and we further calculated that 10^-5^, 10^-4^, and 10^-3^ dilutions of OD600 of 1.0 culture would yield approximately that number of cells per 10 µL injected. *M. sexta* larvae were raised, injected as described above, and then observed over 72 h.

## Results

### Strains ID10 and HGB2511 belong to one of two subspecies of *Xenorhabdus griffiniae* while *Xenorhabdus* sp. TH1 is a novel species

An *X. griffiniae* bacterial isolate (HGB2511) and its nematode host, a strain of *Steinernema hermaphroditum* [24] are being developed as a genetically tractable model for interrogating bacteria-host interactions [24, 25, 76]. To clarify the relationship between HGB2511 and other related *Xenorhabdus* isolates, we sought to comparatively analyse their genomes. We first sequenced the genome of three strains: HGB2511 [24], ID10, the *X. griffiniae* type strain isolated from an Indonesian strain of *S. hermaphroditum* [27], and TH1 isolated from *Steinernema adamsi* from Thailand [77]. For all three strains, we obtained circularised genome assemblies of the bacterial chromosome that had <0.7% contamination, >99.6% completion and >50x depth (Table 1).

We leveraged these high-quality genomes to conduct a wide range of comparative genome analyses, the first of which was taxonomic species delineation via both dDDH and phylogenomics. These analyses both showed the same relationships among the *X. griffiniae* and closely related species (Fig. 1 & Fig. S1 in Additional file 1). The type strain designation of strain ID10 was corroborated by its lack of >70% pairwise dDDH values, the threshold for conspecific strains [78], with any of the other type strains of *Xenorhabdus* (Fig. 1A). Five additional strains were delineated as members of the *X. griffiniae* species, as they each had pairwise dDDH values with strain ID10 that were above the 70% threshold. Among the six *X. griffiniae* strains, two subspecies were evident due to intragroup pairwise dDDH values that were all above the 80% threshold [79]. Strains from Kenya, XN45, VH1, xg97, and Kalro, belonged to one subspecies while strains ID10 and HGB2511 from Indonesia and India, respectively, belonged to another subspecies (Fig. 1A and 1B). Although *Xenorhabdus* sp. TH1 was the most closely related strain to *X. griffiniae* HGB2511 and *Xenorhabdus* sp. BG5, it is an undescribed species of the genus as it lacked pairwise dDDH values, with any of the type strains that were above 70% (Fig. 1A). Indeed, in a pangenome of the seven strains, TH1 had the highest number of strain-specific genes, and none of the ANI pairwise values between strain TH1 and its six closest relatives met the 95% threshold [48] for conspecific strains (Fig. 2A, Supplementary Sheet S13 in Additional file 2). Hence, *Xenorhabdus* sp. TH1 is a novel species of the *Xenorhabdus* genus and not a strain of *X. griffiniae* as was stated in the description of its nematode host [77].

**Fig. 1:**
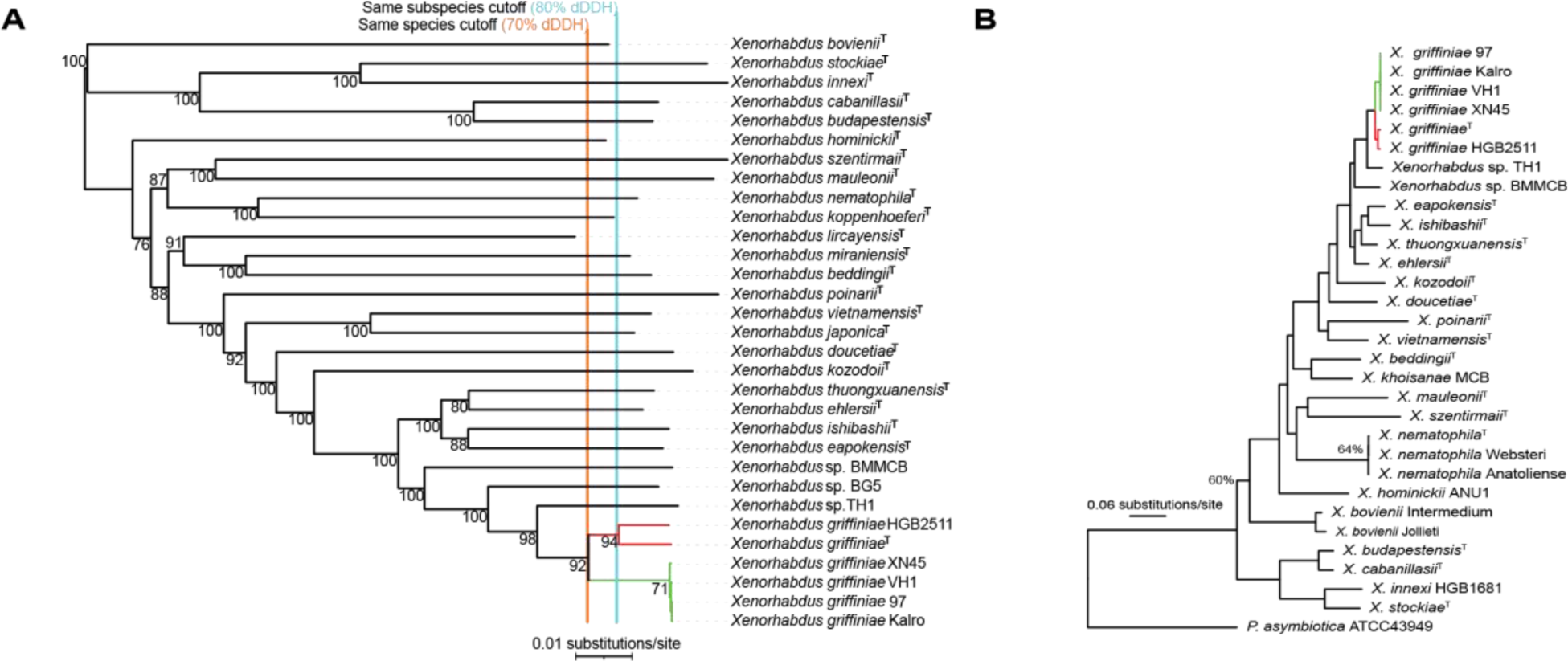
Phylogenomic reconstruction of type strains of *Xenorhabdus* and other strains closely related to *Xenorhabdus griffiniae*. **A)** This neighbour joining tree was reconstructed using genomic distances calculated with the same formula (Genome Distance BLAST Phylogeny distance formula *d5)* used for species delineation by digital DNA-DNA hybridisation (dDDH) analyses. Orange and aqua lines correlate with dDDH boundaries for species (>70%) and subspecies (>80%), respectively. Strains of *X. griffiniae* formed two subspecies, those from India-Indonesia (red) and those from Kenya (green). **B)** Bayesian phylogenetic tree created using one-to-one orthologs from *Xenorhabdus* type strains, *X. griffiniae* strains, and *Photorhabdus asymbiotica* as an outgroup. Posterior probabilities are equal to 1 (100%) unless otherwise indicated at a given node. Strains of *X. griffiniae* formed two separate clades, those isolated from *S. hermaphroditum* nematodes found in Indonesia (ID10) and India (HGB2511) (red) and those isolated from nematodes found in Kenya (green).

**Fig. 2:**
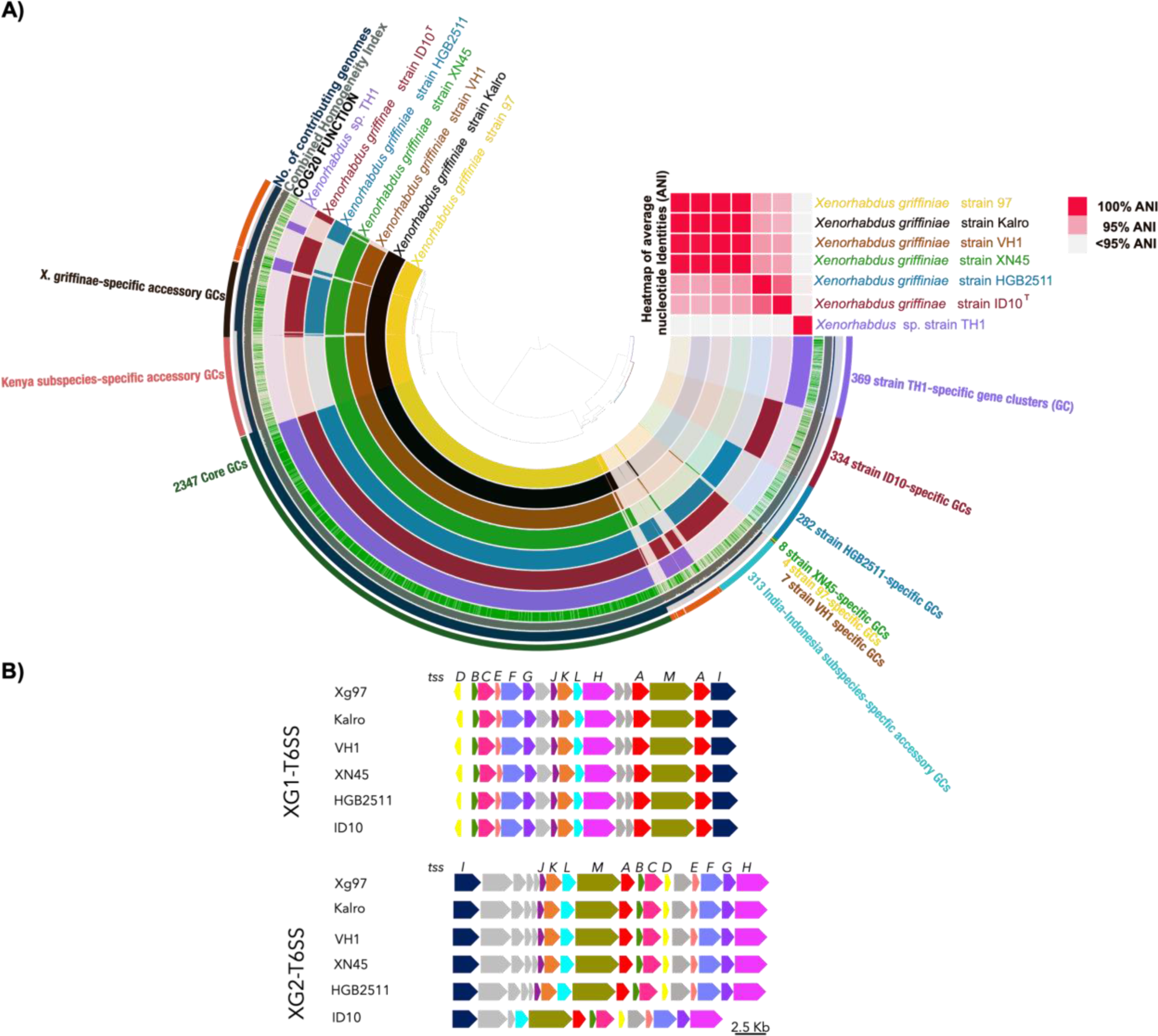
Graphical representations of a pangenome of the *Xenorhabdus griffiniae* clade**. A**) Sunburst chart of a pangenome of the seven closest known relatives of the *X. griffiniae* type strain coupled to a heatmap of their average nucleotide identities (ANI). The pangenome contained 27,337 genes that were clustered into 5,113 groups of orthologs known as gene clusters (GC). The genomes from which each of the GCs was constituted are depicted in the sunburst chart as follows. Each concentric ring represents a genome and each radius represents a GC. For each radius, a dark shade across a concentric ring denotes that the GC is composed of genes from that genome. For the ANI heatmap, shades of red represent pairwise ANI values between 95% (blush) and 100% (rose), the threshold values for conspecific strains. **B)** Loci encoding type six secretions systems (T6SS) found only in *X. griffiniae* genomes, of those analysed here. Each genome encoded two different T6SS, XG1-T6SS and XG2-T6SS. The core T6SS-encoding genes are indicated (*tssA-M*). Other genes are in grey. Pairwise percentage nucleotide identities for shown genomic loci that encode XG1-T6SS and XG2-T6SS ranged between 98-100% and 80-100%, respectively.

### *X. griffiniae* species encode type six secretion systems with subspecies-specific effectors

The pangenome of the six *X. griffiniae* strains plus *Xenorhabdus* sp. TH1 (Fig. 2), the most closely related known species to the *X. griffiniae* clade (Fig. 1), had a total of 27,337 genes. These were grouped into 5113 groups of orthologs, which we termed gene clusters (GC). Out of these, 2347 GCs were core in that they contained orthologs from every genome within the pangenome. On the other hand, a total of 369, 334, 282, 8, 4 and 7 GCs were unique to genomes of TH1, ID10, HGB2511, XN45, xg97 and VH1 respectively. Strain Kalro lacked unique GCs as all its GCs were present in the xg97 genome, even though their respective nematode hosts, *Steinernema* sp. Kalro and *Steinernema* sp. 97 are likely two different undescribed species [80]. Accessory GCs, each of which was composed of orthologs from between two and *n*-1 genomes, were 1739. Among these were 313 and 448 GCs, which were unique to genomes from the India-Indonesia and Kenyan subspecies, respectively. GCs that encode traits that define an *X. griffiniae* strain likely fall among the 319 GCs that were unique to *X. griffiniae* genomes, of which, only 202 (63%) had known functions. Among these, T6SS function was most enriched as it represented 12% of all *X. griffiniae*-specific genes (Sheet S2 in Additional file 2). For the T6SS GCs, those specifically encoding core components *tssA-M* were highly conserved as they had 95-100% combined homogeneity indexes—this is an anvi’o pangenome metric for estimating the similarity of orthologs within a GC calculated from sequence similarities and gap penalties derived from a multiple sequence alignment (MSA) of amino acid sequences [44]. The higher the value the more the positions with identical residues and no gaps within the MSA (Sheet S2 in Additional File 2). Upon deeper investigation, we found that all *X. griffiniae* genomes encode two complete T6SS that we designated XG1-T6SS and XG2-T6SS (Fig. 2B). XG1-T6SS loci were almost identical across six genomes, as the pairwise nucleotide percentage identities for this locus were between 98-100% (Sheet S16 in Additional File 2), and they were found in roughly the same chromosomal location (Fig. 3) in the four circularised genomes (xg97, Kalro, HGB2511 and ID10). For XG2-T6SS, none of the six corresponding genomic loci had pairwise nucleotide sequence identities that were less than 80% (Sheet S16 in Additional File 2). However, the ID10 XG2-T6SS encoding locus uniquely lacked *tssK, tssJ* and two other genes that were directly downstream of *tssJ* (Fig. 2). Like XG1-T6SS, the XG2-T6SS loci were also found in a similar chromosomal region across the four circularised genomes (Fig. 3). Based on the high pairwise nucleotide identities, we identified homologs of XG1-T6SS and XG2-T6SS-encoding loci in *X. szentirmaii* US123, *X. doucetiae***^T^**, *X. cabanillasi***^T^**, *X. hominickii* ANU, *X. nematophila***^T^**, *X. poinarii***^T^** and *X. bovienii* SS-2004 (Sheet S16 in Additional File 2). The strain SS-2004 homologs were those identified by Chaston *et al.* [81] and designated T6SS-1 and T6SS-2, respectively, by Kochanowsky *et al.* [82].

**Fig. 3:**
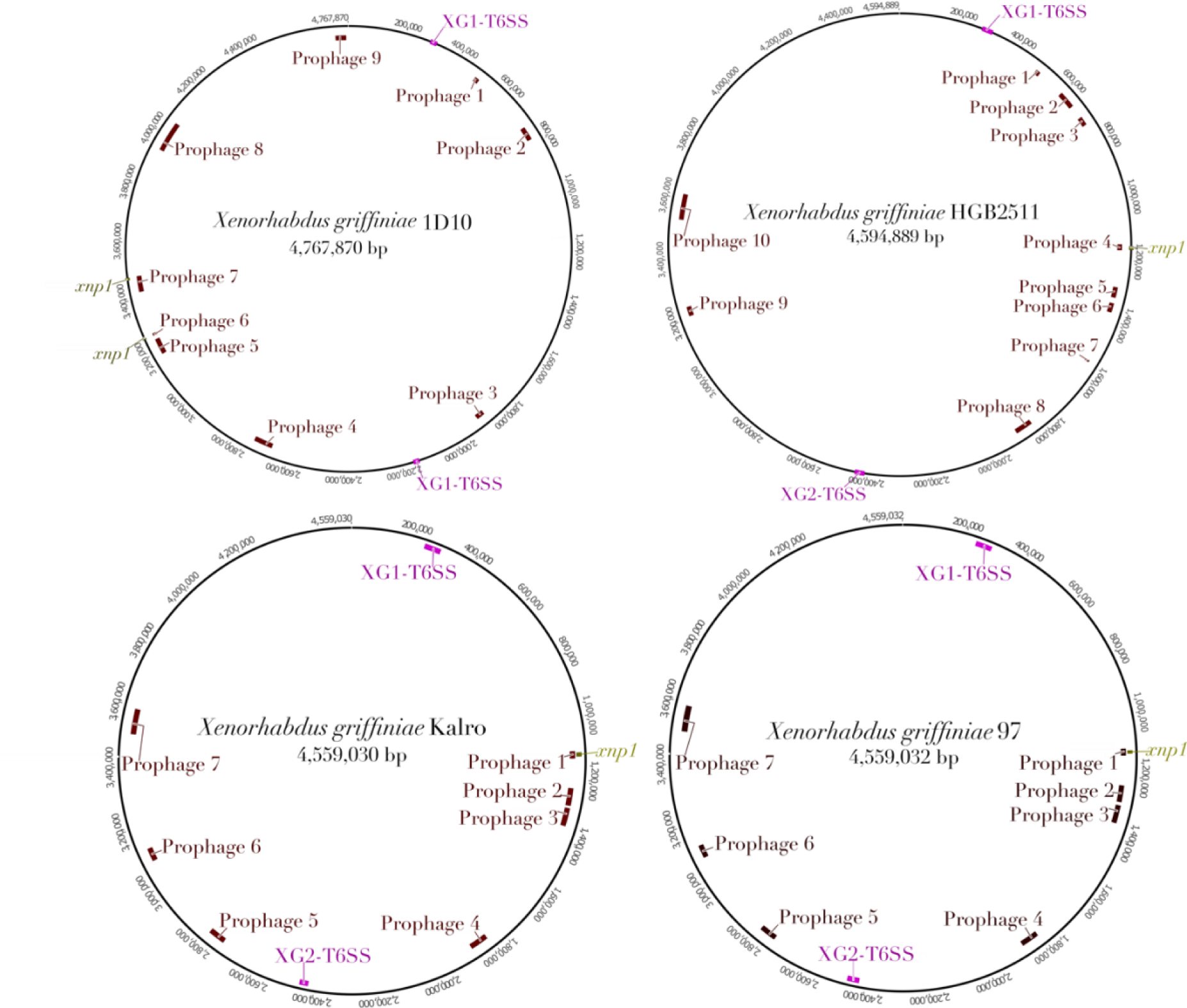
Loci of prophages and gene clusters encoding complete type six secretion systems (T6SS) and xenorhabdicin (*xnp1*) in complete genomes of four *Xenorhabdus griffiniae* strains.

We further identified T6SS-associated GCs that are subspecies-specific. The Kenyan and India-Indonesia subspecies have eight and nine subspecies-specific GCs, respectively. The majority of these are predicted to encode spike proteins annotated as VgrG or PAAR-domain-containing Rhs proteins (Sheet S3 in Additional File 2). One Kenyan subspecies GC encodes a *tssF* that was not part of the two complete T6SS-encoding loci. We analysed the genes in the neighbourhood of the subspecies-specific PAAR-encoding loci for genes that encode T6SS effector proteins and their cognate immunity proteins. We found four such loci that are specific to the India-Indonesia subspecies and that share similar gene content and synteny (Fig. S4 in Additional File 1). For each of these four loci, their encoded PAAR proteins are highly similar since their amino acid sequences had combined homogeneity indexes between 94-100% (Sheet S3 in Additional File 2). These findings indicate that in *X. griffiniae,* the T6SS spike and its cognate effector proteins may contribute to intraspecific traits. We extended these findings by predicting other putative secretion systems encoded across the sequenced genomes using TXSScan (Table S1 in Additional File 1). The number of type I systems varied across the genomes analysed, and at least one copy of flagellum, Type 4a pilus (T4aP), Type 5a Secretion system (T5aSS), and Type 5b secretion system (T5bSS) were identified in all the genomes analysed.

### Prophages mediated the acquisition of both subspecies-specific and strain-specific genes

We hypothesized that horizontal gene transfer was a major driver of subspeciation in *X. griffiniae*, since the mobilome constituted the largest fraction of functionally annotated, strain-specific genes (Fig. 4, sky blue=mobilome: prophages, transposons, plasmids), and strain-specific genes result in speciation when they confer ecologically useful traits [83].

**Fig. 4:**
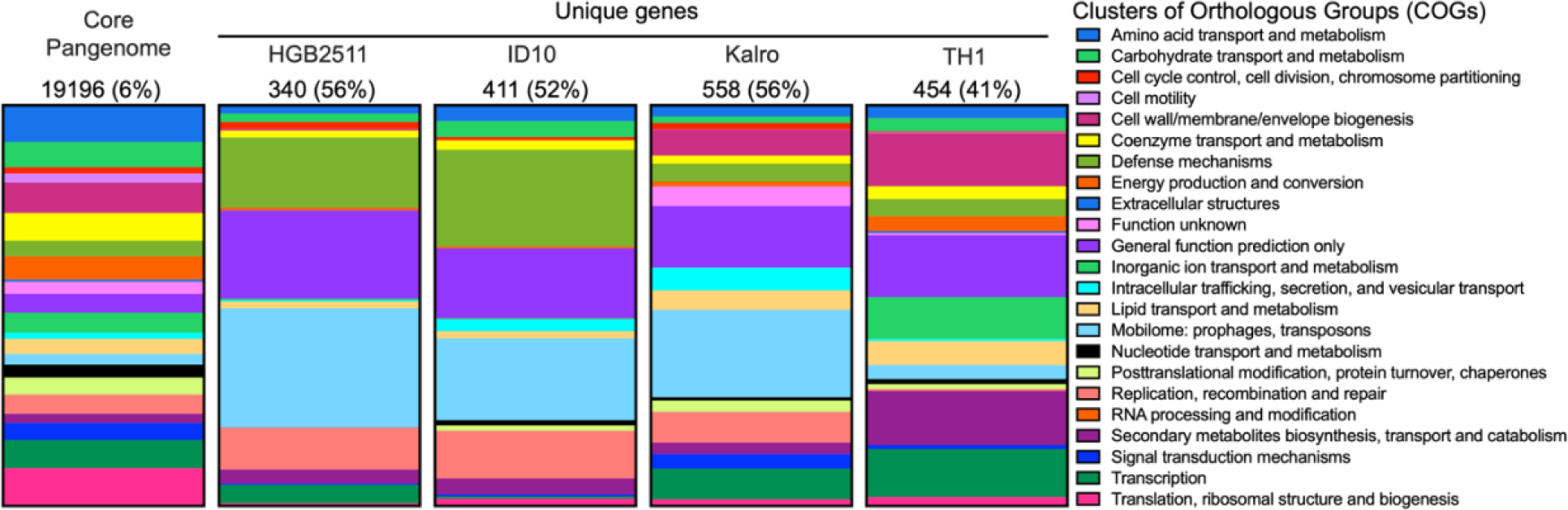
Stacked bar charts depicting the Clusters of Orthologs Groups of proteins (COGs) of the core and unique genes among four closely related *Xenorhabdus* strains. The core pangenome bar shows the COGs of genes that are common to all genomes in the pangenome. The subsequent bars show the COGs of the unique genes from the HGB2511, ID10, Kalro, and TH1 genomes derived from the previously described pangenome analysis of *X. griffiniae* strains and *Xenorhabdus* sp. TH1 (Fig. 2A). The numbers on top of each bar are the total number of genes in each category, followed by the percentage of those genes without COG designation in parentheses.

We first investigated this by inferring the evolution of gene content among the six strains of *X. griffiniae* and *Xenorhabdus* sp. TH1. We inferred that a net gene loss resulted in the speciation of *Xenorhabdus* sp. TH1 (Fig. 5), consistent with its smaller genome size (Table 1).

**Fig. 5.**
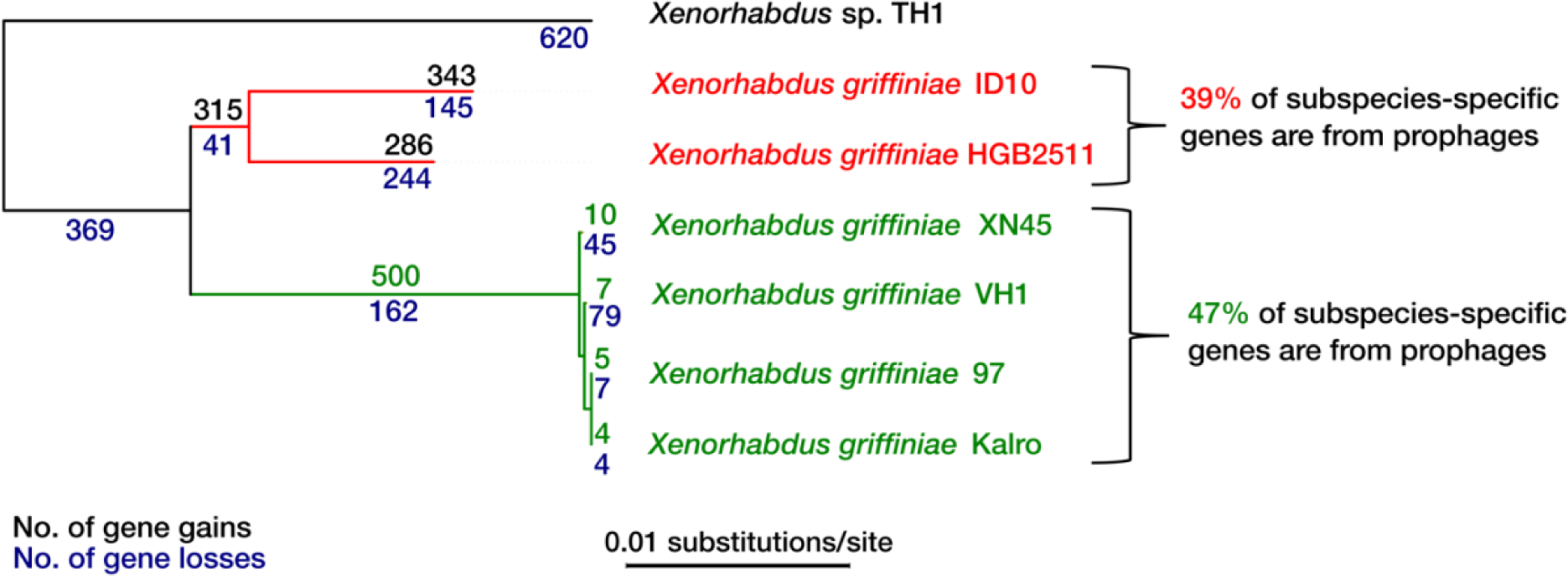
Neighbour-joining phylogenomic tree depicting the evolution of gene content among strains of *Xenorhabdus griffiniae*. Two species are depicted in this tree, *Xenorhabdus* sp. TH1 and *X. griffiniae*. For *X. griffiniae,* its India-Indonesia and Kenya subspecies are in red and green, respectively. The emergence of both subspecies was due to net gene gains. Gene content analysis was conducted in COUNT applying Wagner parsimony.

Conversely, net gene gains resulted in the formation of the two *X. griffiniae* subspecies. We addressed the question of whether horizontal gene transfer (HGT) may have mediated these gene gains by conducting a preliminary pangenome analysis of 49 *Xenorhabdus* strains. We found that the total number of phage-related genes accounted for 55% of the variation in the proteome sizes among *Xenorhabdus* genomes. In this analysis, the total number of phage-related genes accounted for 48.69% of the variation in proteome sizes among the seven strains (adjusted r^2^*=*0.48691, *p*=0.04899983). Similar correlations for transposable elements and plasmid-related genes were insignificant (adjusted r^2^*=*0.23241, *p*=0.15, adjusted r^2^*=* 0.1762, *p*=0.19, respectively). Based on this, we focused on the identification of subspecies-specific and strain-specific genes that were linked with prophages.

We first identified prophages in TH1, ID10, HGB2511, Kalro and xg97 (Fig. 3 and Fig. S2 in Additional File 1, Sheets S8-S11 and S15 in Additional File 2). Genomes of VH1 and XN45 were excluded as they were too fragmented to yield robust results. Taxonomically, all identified prophages in the five genomes belonged to the family *Caudoviricetes* (Sheet S15 in Additional File 2). The genomes of HGB2511, ID10, Kalro, 97 and TH1, had ten, nine, seven, seven, and three prophages respectively.

To determine how these prophage numbers compared to those found in other strains, we similarly identified prophages in ten other *Xenorhabdus* strains whose chromosomal genomes were also assembled into one contig (Sheet S7 in Additional File 2). We found an average of seven prophages per genome, ranging from 3 to 13, indicating that HGB2511 and ID10 harbour higher-than-average numbers of prophages in their genomes. Three prophage loci were similar across genomes: 1) ID10 prophage 3 and HGB2511 prophage 3 which had 73% pairwise nucleotide percentage identities; 2) ID10 prophage 9 and HGB2511 prophage 5 with 61% pairwise nucleotide percentage identities (Fig. S3 in Additional File 1); and 3) the locus (*xnp1*) [84], whose conserved and variable regions were previously elucidated in strains of *X. nematophila* and *X. bovienii* and shown to encode xenorhabdicin, an antimicrobial R-type pyocin, or tailocin structure [85]. The *xnp1* locus, including genes essential for xenorhabdicin production and release, were detected in all seven genomes (Fig. 3; Fig. 6; Fig. S2 in Additional File 1).

**Fig. 6.**
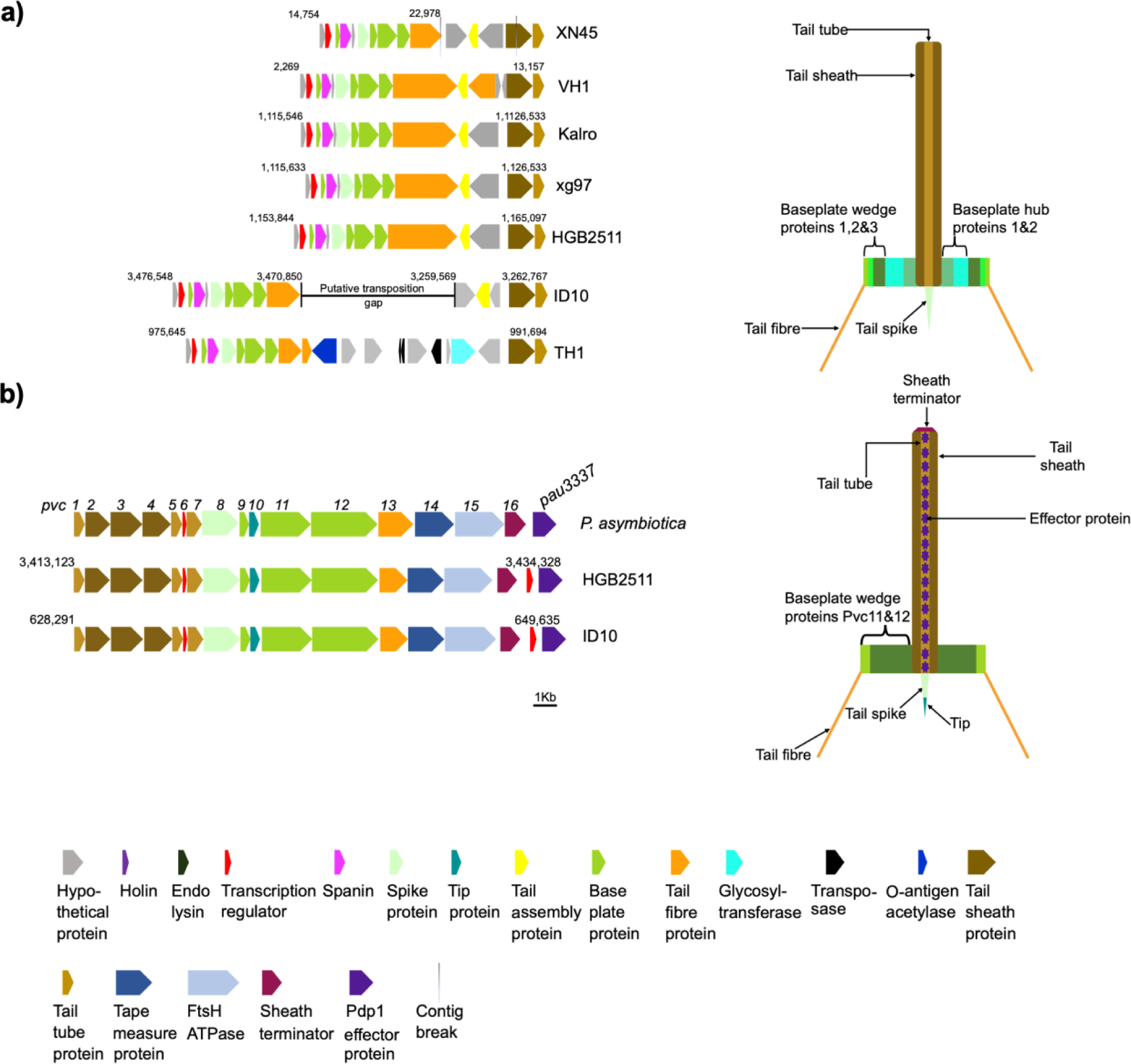
Genomic loci encoding phage tail-like particles from *Xenorhabdus griffiniae.* A) The xenorhabdicin encoding genomic loci (*xnp1*) in six strains of *Xenorhabdus griffiniae* and *Xenorhabdus* sp. TH1 and a cartoon of the corresponding xenorhabdicin particle. B) Genomic loci (*pvc*) encoding an extracellular contractile injection system, the *Photorhabdus* Virulence Cassette (PVC) in *P. asymbiotica* ATCC43949, *X. griffiniae* ID10 and HGB2511. The genome coordinates of the loci are shown for the circularised genome assemblies. For XN45 and VH1, the shown coordinates are for contigs JACWFC010000129 and JADEUF010000065, respectively.

Comparative analysis of *xnp1* loci revealed that in ID10 it has been split into two loci that are two megabase pairs apart (Fig. 3 & Fig. 6), probably due to transposition events. The TH1 *xnp1* locus included genes that encode O-antigen acetylase and glycosyltransferases (Fig. 6) which may be involved in conferring immunity to xenorhabdicins through modification of its likely receptor, lipopolysaccharide O antigen [86].

To identify the specific genes contained in the identified prophage regions, we took ID10 prophages as an example (Fig. 7). Genes predicted to encode viral replication and hypothetical proteins constituted 35 and 15 percent, respectively (Sheet S19 in Additional File 2) whereas ‘cargo’ genes with non-virus, functional annotations constituted the remaining half. These annotated cargo genes encoded diverse products, including toxin-antitoxin systems that have wide-ranging effects on bacterial physiology and mobile genetic elements within genomes [87], Importin-11, predicted to encode a nuclear transport receptor that presumably would be delivered for modulation of animal host cell physiology [88], and diguanylate cyclase, predicted to be part of a signal transduction cascade mediated through the second messenger cyclic-di-GMP [89] (Fig. 7).

**Fig. 7:**
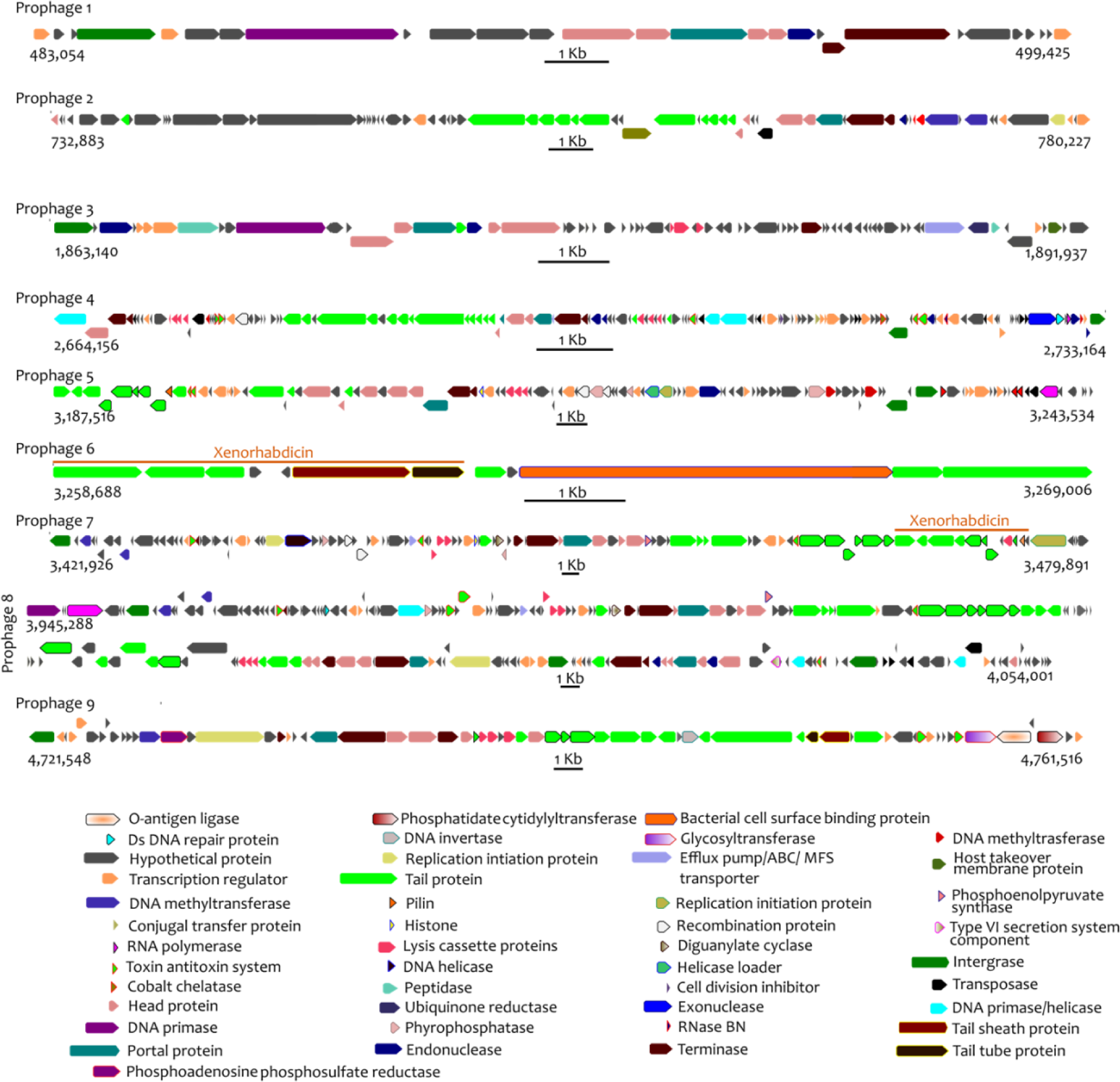
Graphical representation of predicted products of genes contained within prophage loci of *Xenorhabdus griffiniae* ID10. The genes could be categorized into three broad categories; annotated cargo genes, viral replication genes and those whose products are unknown.

The identified prophages contained 45 and 38% of strain-specific genes in the HGB2511 and ID10 genomes, respectively. Likewise, 39 and 47% of India-Indonesia and Kenyan subspecies-specific genes were from prophages (Fig. 5). Indeed, removal of prophage regions from the four genomes elevated their pairwise dDDH values: Pairwise dDDH values between ID10 and strains HGB2511, Kalro, TH1 and xg97 rose by 3-3.2 percentage points when identified prophages were removed from all genomes (Sheet S12 in Additional File 2). These findings demonstrate that, in *X. griffiniae*, a considerable proportion of both subspecies and strain-specific genes were gained through prophages.

### Comparative analysis of CRISPR-Cas, protospacer, and anti-CRISPR content

The high prevalence of prophages and prophage-mediated gene gains in *X. griffiniae* genomes suggests that these bacterial symbionts relatively frequently encounter phage-related foreign DNA. This prompted us to investigate the presence or absence of defence systems such as the CRISPR-Cas immunity system [90]. We found that the genomes of each of the analysed *X. griffiniae* genomes and the close relative TH1 encode a syntenic locus containing a full set of Class1-Subtype-I-E *cas* genes encoded adjancet to a *gntR* homolog (Fig. 8A). The former closest known relative of ID10, BMMCB, has an incomplete set of *cas* genes on a single contig, with *casD* and *cas2* lacking. However, since the BMMCB genome is fragmented, we cannot rule out the possibility that these genes are encoded elsewhere. HGB2511, ID10, and TH1 also have a second set of Class1-Subtype-I-E genes (but lacking *cas1* and *cas2*) encoded adjacent to an *eda* homolog (Fig. 8B). All strains except ID10 and BMMCB also have three CRISPR arrays, comprising conserved repeats and variable targeting spacers that are predicted to be transcribed and cleaved into non-coding, small (61 nt), targeting CRISPR RNAs (crRNAs): array 1a, adjacent to the full set of *cas* genes at the *gntR* locus (Fig. 8A,D), array 1b, adjacent to the *eda* homolog (Fig. 8B,E), and array 2, adjacent to a *dsbB* gene (Fig. 8C,F). BMMCB lacks the CRISPR arrays adjacent to *eda* and *dsbB*. Instead, BMMCB has a second CRISPR array with just two spacers in another region of the genome (at coordinate 3720455, not shown in figure) that encodes phage-related genes. This array falls at the edge of a contig break, so may not be an accurate reflection of the repeats that might be present at this locus.

**Fig. 8:**
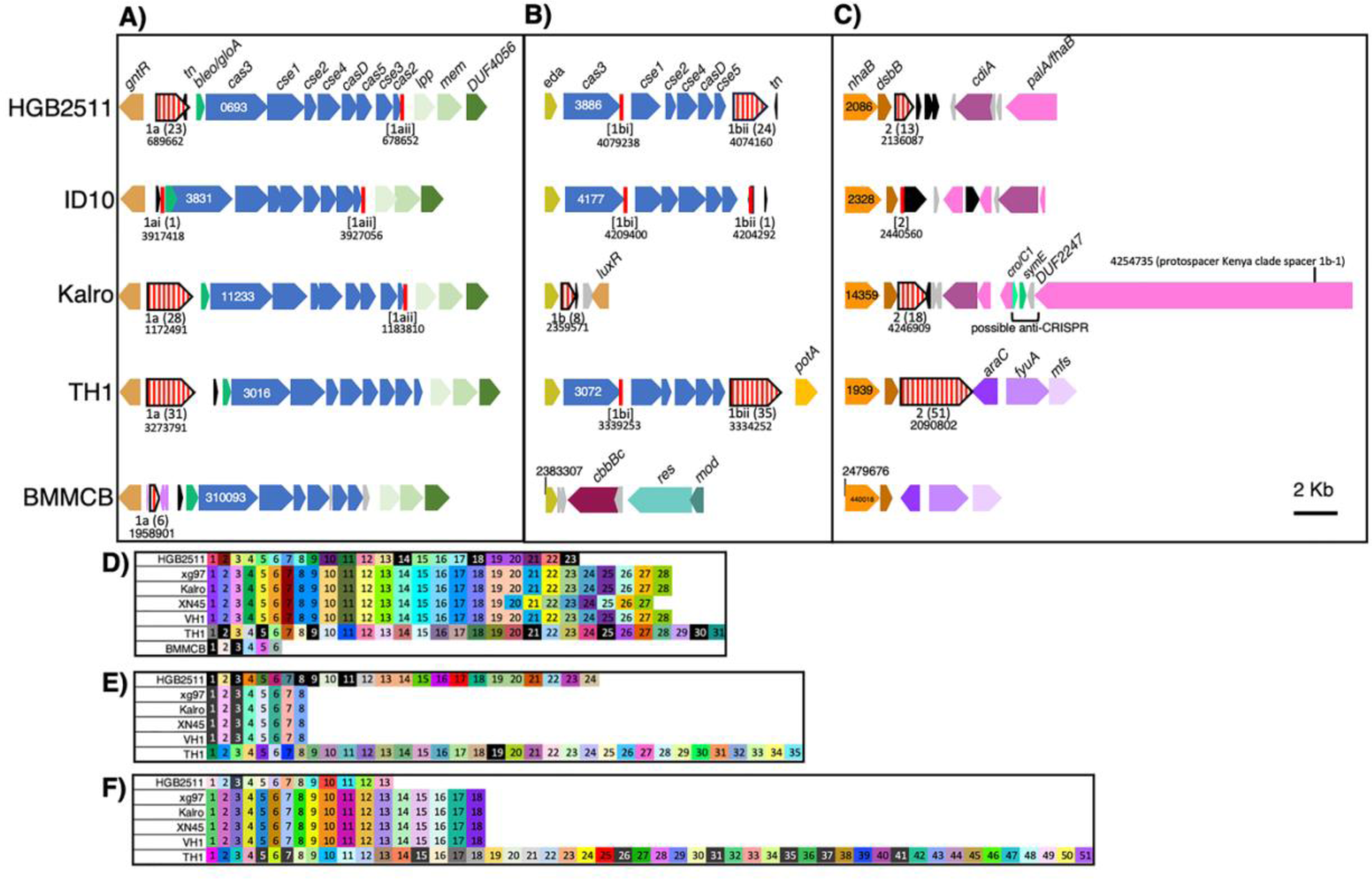
CRISPR-Cas regions of *Xenorhabdus* strains. **A-C)** Comparison of three CRISPR-Cas related genomic regions in selected strains, anchored for synteny in the diagram using *gntR* **(A)**, *eda* **(B)**, or *nhaB* **(C)**. ORFs are indicated by solid block arrows (blue for *cas* genes, with the locus tag number of *cas3* provided) with annotated gene names indicated above, and identical colours indicating homology. CRISPR repeat arrays (red vertical stripe block arrows) for each region (1a, 1b, 2) had variable number of repeats (noted in parentheses). Start coordinates for each are shown underneath. Degenerate repeat arrays are indicated by brackets. A conserved sequence [1aii; vertical red line] containing a repeat, a spacer, and a degenerate repeat) was identified at the end of *cas2,* in all strains except BMMCB, which lacks this gene **(A)**. Another [1bi] was apparent at the end of *cas3* in the *eda* locus of HGB2511, ID10, and TH1 **(B)**. **D-F)** The spacer sequences of each CRISPR array found at *gntR* **(D)**, *eda* **(E),** or *nhaB* **(F**) were compared for identity to each other or other loci among the analysed strains. Each box represents a spacer, and different colours indicate different sequences. Spacers represented by black boxes and white lettering have 100% identity to “target” loci outside of the array, either within the same genome or within one of the other genomes analysed here. *gntR:* DNA-binding transcriptional repressor; *bleo/gloA*: bleomycin resistance/glyoxalase; *tn*: transposase; *lpp:* lipoprotein; *mem*: membrane protein: DUF4056: domain of unknown function 4056 gene; *eda:* 4-hydroxy-2-oxoglutarate aldolase; *nhaB*: Na(+):H(+) antiporter NhaB; *dsbB:* protein thiolquinone oxidoreductase; *cdiA*: Deoxyribonuclease CdiA; *palA/fhbA*: filamentous hemagglutinin; *luxR*: LuxR family transcriptional regulator; *cro/CI*: HTH cro/C1-type domain-containing protein; *symE*: Type I addiction module toxin, SymE family; *potA*: spermidine preferential ABC transporter ATP binding subunit; *araC:* AraC family transcriptional regulator; *fyuA:* Putative TonB-dependent siderophore receptor; *mfs*: putative MFS transporter, signal transducer; *cbbBC*: Molybdopterin-binding oxidoreductase; *res*: Type III restriction endonuclease subunit R; *mod*: site-specific DNA-methyltransferase (adenine-specific).

Since spacer sequences are acquired in a directional manner in response to active infection by foreign nucleic acid material (e.g., phages or plasmids), comparisons of spacer content across related strains can be used to infer their shared life histories and prior exposure to such threats [91, 92]. The spacer contents of CRISPR arrays 1a, 1b and 2, is variable in number and sequence across the strains (Fig. 8D-F). Consistent with the close relatedness of the Kenyan subspecies strains xg97, Kalro, XN45, and VH1, their CRISPR spacer content is identical, except for the absence in XN45 of a duplicated spacer found in array 1a of the other strains (Fig. 8D). Otherwise, there is no overlap in spacer identity among CRISPR arrays of the different strains, indicating their divergence prior to the acquisition of existing spacer content. Compared to the other *X. griffiniae* and the close relative TH1, the CRISPR arrays in ID10 appear to have a limited number of targeting spacers. Based on the presence of conserved repeat sequences, ID10 encodes two (1ai and 1bii) bona fide, single repeat CRISPR loci at locations syntenic with regions 1a and 1b of the other strains. Three other loci (1aii, 1bi, and 2) appear to be remnants, with only a single clear left repeat, a spacer, and a degenerate right repeat (Fig. 8A-C; Additional File 3).

To gain insights into the types of threats encountered by *X. griffiniae* and related strains, we searched for putative target sequences (known as protospacers) based on their identity with CRISPR array spacer sequences. We found some spacers have 100% identity to protospacers either within the same genome (self-targeting) or within one of the other genomes analysed here (Table 2; Table S5 in Additional File 3). In many cases, these protospacers were within phage-related, conjugation machinery, and restriction modification systems, in line with the role of CRISPR systems in defending against these types of mobile genetic elements [93, 94]. Consistent with their close relationship, the Kenya clade demonstrated identical protospacer content in genes throughout the genome, including several predicted to be targeted by CRISPR small RNAs from HGB2511 (Table 2; Table S5 in Additional File 3).

Additional functional genes with protospacer sequences that could be targeted by crRNA included those predicted to encode filamentous hemagglutinin (Kenya clade spacer 1b-1), an ABC transporter (Kenya clade spacer 1b-3), and the enzymes FolD (TH1 spacer 1a-5) and GcvP (BMMCB spacer 1a-3). Curiously, the *palA/fhaB* filamentous hemagglutinin gene with self-identity to the Kenya clade spacer 1b-1 is encoded in the *dsb* locus, in proximity to the Kenya clade spacer region 2 (Fig. 8C). Since spacer self-identity would presumably result in self-intoxication, we hypothesize the genome also encodes an anti-CRISPR immunity mechanism such as anti-CRISPR proteins known as Acr. These proteins are difficult to predict with sequence similarity because they vary widely [94]. We manually searched for such loci in the selected genomes using a “guilt-by-association” approach of putative Acr by identifying small open reading frames in proximity to the protospacer-containing gene and a helix-turn-helix (HTH) domain containing gene, which is predicted to be the anti-CRISPR regulator. Of the self-targeting protospacers we detected, only the one in *palA/fhaB* of the Kenyan subclade had a promising candidate based on these criteria (Fig. 8C; Additional File 3). The putative Acr is a DUF2247 domain-containing protein (e.g., JASDYB01_14371) which is predicted to encode a protein of 171 aa and is encoded near an HTH cro/C1-type domain-containing protein (e.g., JASDYB01_14369) that may be a putative Aca transcriptional regulator [94].

### Restriction Modification Systems

In addition to CRISPR arrays, restriction-modification systems resist the introduction of foreign DNA, including phage infection, by detecting and cleaving non-chromosomal DNA. Restriction-modification systems can be classified based on their structure, cofactor requirements, DNA recognition site and relative cleavage locations [95, 96]. Type I, II, and III all encode both a restriction endonuclease and a methyltransferase, whereas Type IV endonucleases cleave modified DNA (such as 5-hydroxymethylcytosine) and variably encode an adjoining methyltransferase [96]. There are also anti-restriction proteins which inhibit restriction modification systems by various mechanisms. We predicted the number and type of complete restriction-modification systems and anti-restriction proteins in the BMMCB, TH1, HGB2511, ID10 and Kalro genomes (Table S2 in Additional File 3). Type I and II systems were the most prevalent across the genomes. Only the HGB2511 genome encoded a complete Type III system, while ID10, Kalro, and TH1 genomes each encoded Type IV restriction endonucleases. The genomes of all *X. griffiniae* and *Xenorhabdus* sp. TH1 encoded at least one anti-restriction protein, with ID10 appearing to encode seven anti-restriction proteins, by far the largest complement (Table S2 in Additional File 3).

### *X. griffiniae* encode the biosynthesis of diverse natural products

*Xenorhabdus* genomes often comprise loci, known generally as biosynthetic gene clusters (BGCs), which are responsible for the biosynthesis of a wide array of natural products. A BGC can include many genes, often under the control of one promoter, which collectively encode the production pathway of a single natural product and its derivatives. Commensurate with its genus, which ranks among those that produce the most diverse set of natural products [97], the *X. griffiniae* ID10 genome contained over 21 biosynthetic gene clusters that were predicted to encode the production of over ten different types of natural products (Fig. 9; Table S3 in Additional File 1).

**Fig. 9:**
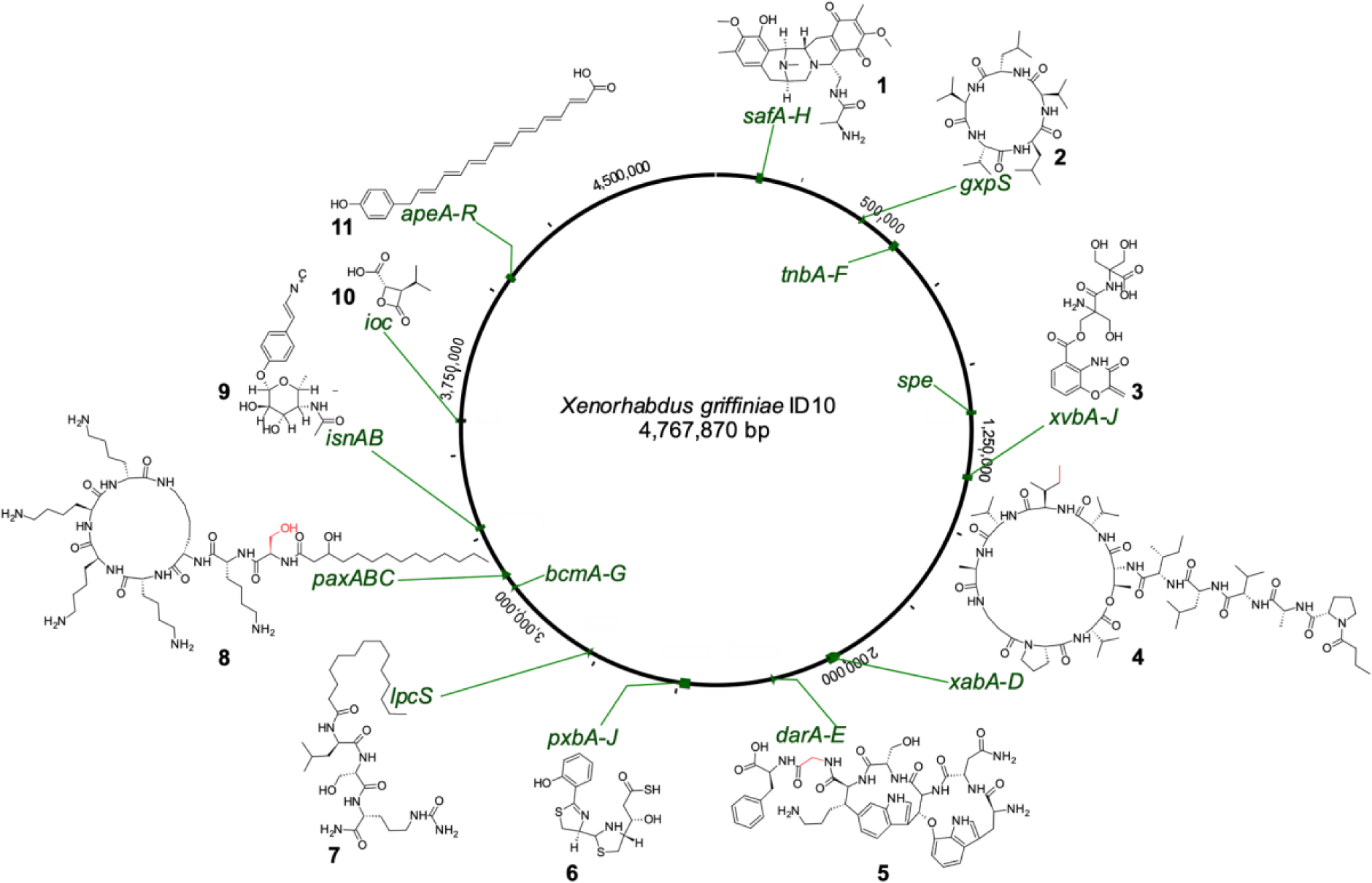
Genomic loci of known biosynthetic gene clusters in the *X. griffiniae* ID10 genome and predicted chemical structures of the natural products whose biosynthesis they encode. The *paxABC*, *darABCDE* and *xabABCD* BGCs were predicted to encode the production of potentially novel derivatives of PAX peptides (8), darobactin (5) and xenoamicin (4), respectively, that differed from known structures in amino acid building blocks at positions highlighted in red. Safracin (1), gameXpeptide C (2), benzobactin (3), photoxenobactin (6), type 2 bovienimide (7), rhabduscin (9), 3-isopropyl-4-oxo-2-oxetanecarboxylic acid (10), arylpolyene (11).

Fourteen of these ID10 biosynthetic gene clusters (BGCs) are predicted to encode the biosynthesis of either known compounds or their derivatives (Fig. 9; Table S3 in Additional File 1). For example, the *lpcS* and *isnAB,GT* BGCs in the ID10 genome are predicted to encode the production of group IIA bovienimides [98] and rhabduscin [99], respectively, both of which are insect immunity suppressors, the *pxb* BGC which encodes the production of the insecticidal photoxenobactins [20], and *safA-H, ioc/leu, xvbA-J, bcmA-G,* and *ape* BGCs which respectively encode the production of safracin antibiotics, 3-isopropyl-4-oxo-2-oxetanecarboxylic acid (IOC), benzobactins, bicyclomycins, and aryl polyenes [20, 100–102]. The *gxpS* BGC was predicted to encode the synthesis of GameXPeptide C [103], as their predicted peptide sequence was ^D^Val-^L^Val-^D^Leu-^L^Val- ^L^Leu. In contrast, each member of the *X. griffiniae* India-Indonesia subspecies had a unique set of BGCs. Specifically, the HGB2511 strain lacked BGCs that encoded the production of rhabduscin, benzobactin, bicyclomycin and actinospectacin, all of which were present in the ID10 genome (Table S3 in Additional File 1).

Notably, the ID10 genome contained known BGCs but the predicted biosynthetic products are previously unknown derivatives. For example, the ID10 *paxABC* BGC, which encodes the biosynthesis of PAX peptides, is predicted to encode an octapeptide backbone of ^L^Ser-^L^Lys-^L^Lys-^D^Lys-^D^Lys-^D^Lys-^D^Lys, which differs from those of *X. nematophila* [104] and *X. khoisanae* [105] by having ^L^Ser at position one instead of Gly, since the respective Stachelhaus code was DVWHLSLIDK and not DILQIGLIWK. The *xabABCD* BGC is predicted to encode the biosynthesis novel xenoamicins that incorporate ^D^Ile in lieu of ^D^Val [106] at position eight of the tetradodecapeptide backbone. The synthesis of novel derivatives is also predicted for BGCs that encode the biosynthesis of the known ribosomally-synthesized and post-translationally modified peptide (RiPP), darobactin [107], since the ID10 *darABCDE* BGC was predicted to encode the biosynthesis a core peptide with the sequence Trp-Asn-Trp-Ser-Lys-Gly-Phe and not Trp-Asn-Trp-Ser-Lys-Ser-Phe.

### *X. griffiniae* encode entomotoxins and are insecticidal to *Manduca sexta*

An essential part of the *Steinernema-Xenorhabdus* entomopathogenic lifecycle, is the ability of the host-symbiont pair to infect and kill insect hosts, and *Xenorhabdus* produce virulence factors that target other microorganisms competing for the nutritious insect cadaver [7]. To better understand the toxic potential of the *X. griffiniae* bacteria, we mined the HGB2511, ID10, Kalro and TH1 genomes for toxin-domain-containing loci. Using a list of known toxins found in other *Xenorhabdus* [19, 68] we identified homologs of genes encoding the known insecticidal toxins Mcf “makes caterpillars floppy” and PirAB in each of the strains [108–110], along with proteins homologous to the MARTX toxin family (Table 3). Similar to *X. innexi* HGB1681, the MARTX proteins in HGB2511, ID10, Kalro and TH1 each lack four of the A repeats at the *N*-terminus of the protein (A Δ3-7), leaving nine repeats compared to the 14 found in *X. nematophila* 19061, *X. bovienii* SS-2004 (Jollieti), and *Vibrio* species [68, 111] (Fig. S5 in Additional File 1). It remains unclear how these differences in repeat structure might impact MARTX protein function in *X. griffiniae*.

Insecticidal Tc toxins are three-part toxin complexes (TcA, TcB, and TcC type) commonly found in entomopathogenic bacteria, including in some *Xenorhabdus* species [112]. Tc toxin family proteins were notably absent from the *X. griffiniae* HGB2511, ID10 and Kalro genomes and the *Xenorhabdus* sp. TH1 genome (Table 3). No evidence was found of homologs of Shiga toxin (*stx1a*) related genes, such as those found in *X. bovienii* (Table 3) [19]. Within the ID10 genome we identified a putative insecticidal toxin (XGHID_v1_0629), a homolog of which was also found in the HGB2511 genome (XGHIN1_v1_3228). These genes are homologs of a *Photorhabdus asymbiotica* gene (PAU_03337) (Fig. 6B) that encodes *Photorhabdus* dNTP pyrophosphatase 1 (Pdp1), a cytotoxic protein that not only kills immune cells by reducing their intracellular deposits of deoxynucleotide triphosphates (dNTPs) but is also an effector protein of the extracellular contractile injection system known as *Photorhabdus* virulence cassette (PVC) [113]. Indeed, analysis of genes upstream *pdp1* revealed that both ID10 and HGB2511 encode PVCs (Fig 6B). Although PVCs are phage tail-like particles that are structurally similar to xenorhabdicin (Fig 6B), they differ by having within their tube, effector proteins that are translocated into the target cell, upon tail fibre-mediated binding and subsequent tail sheath contraction [114]. The *N*-terminus (50aa) of the *Photorhabdus asymbiotica* Pdp1 acts as a signal peptide for secretion through the PVC [115]. Amino acid alignment of the ID10 and HGB2511 Pdp1 proteins with the *Photorhabdus* Pdp1 and two non-PVC secreted homologs [115] revealed amino acids at the *N*-terminus of the *Xenorhabdus* proteins that could act as a signal peptide for PVC secretion (Fig. S6, Additional File 1).

A *de novo* search for other toxin homologs within our genomes of interest revealed two strain-specific loci of particular interest [69]. The *Xenorhabdus* sp. TH1 genome contains a complete hydrogen cyanide synthase locus (*hcnABC)* (Table 3). *hcnABC* is found in plant-associated and entomopathogenic bacteria [116] where it plays a role in insect killing. Notably, *hcnABC* was recently identified in the genome of a steinernematid-associated *Pseudomonas piscis* bacterium [80]. In the *X. griffiniae* Kalro genome, three proteins with zonular occludens toxin (zot) domains were identified (JASDYB01_14222, JASDYB01_14237, JASDYB01_14239) (Table 3). Zonula occludens toxin (Zot) domain-containing proteins target the eukaryotic cell cytoskeleton and compromise the structure of intercellular tight junctions, leading to a permeabilization of epithelia [117, 118]. Homologs of the three Zot domain-containing proteins found in Kalro were also identified in xg97, VH1 and XN45, and BLASTp revealed other Zot domain-containing proteins in other *Xenorhabdus* species (including *X. bovienii, X. khoisanae, X. eapokensis, X. ehlersii* and *X. innexi*). JASDYB01_14222 and JASDYB01_14237 are each predicted to encode a transmembrane helix and to be membrane embedded, whereas JASDYB01_14239 is a considerably shorter peptide lacking both transmembrane domains and secretory signals. These Zot domain-containing proteins may affect the insect midgut as part of an oral route of infection [119], or destroy insect epithelial tissues when the bacteria are released into the hemocoel.

The diversity of toxin coding potential within the analysed *Xenorhabuds* genomes suggest possible differences in their entomopathogenicity. To begin to interrogate this possibility, we assessed the survival of fifth instar *Manduca sexta* insect larvae over a 72 h period after injection with five *Xenorhabdus* strains at three concentrations. We aimed to compare the strains at an inoculum of ∼1000 cells because at that dosage our controls, *X. nematophila* (19061) and *X. innexi* (HGB1681) were previously shown to induce near to 100% lethality and <10% lethality, respectively [68], and indeed, these trends were recapitulated in our study (Fig. 10). Each bacterial strain displayed a dose-dependent survival response, with the highest inoculum resulting in the greatest mortality (Fig. S7, Additional File 1). Insects injected with ∼1000 cells of *X. griffiniae* ID10 displayed robust survival, like *X. innexi,* whereas greater than 50% of the animals injected with approximately the same number of *X. griffiniae* HGB2511 and *Xenorhabdus* sp. TH1 cells succumbed within 72 h of injection, like the level observed for *X. nematophila* (Fig. 10). These results reveal that the isolates tested have different levels of virulence against Lepidopteran insects.

**Fig. 10:**
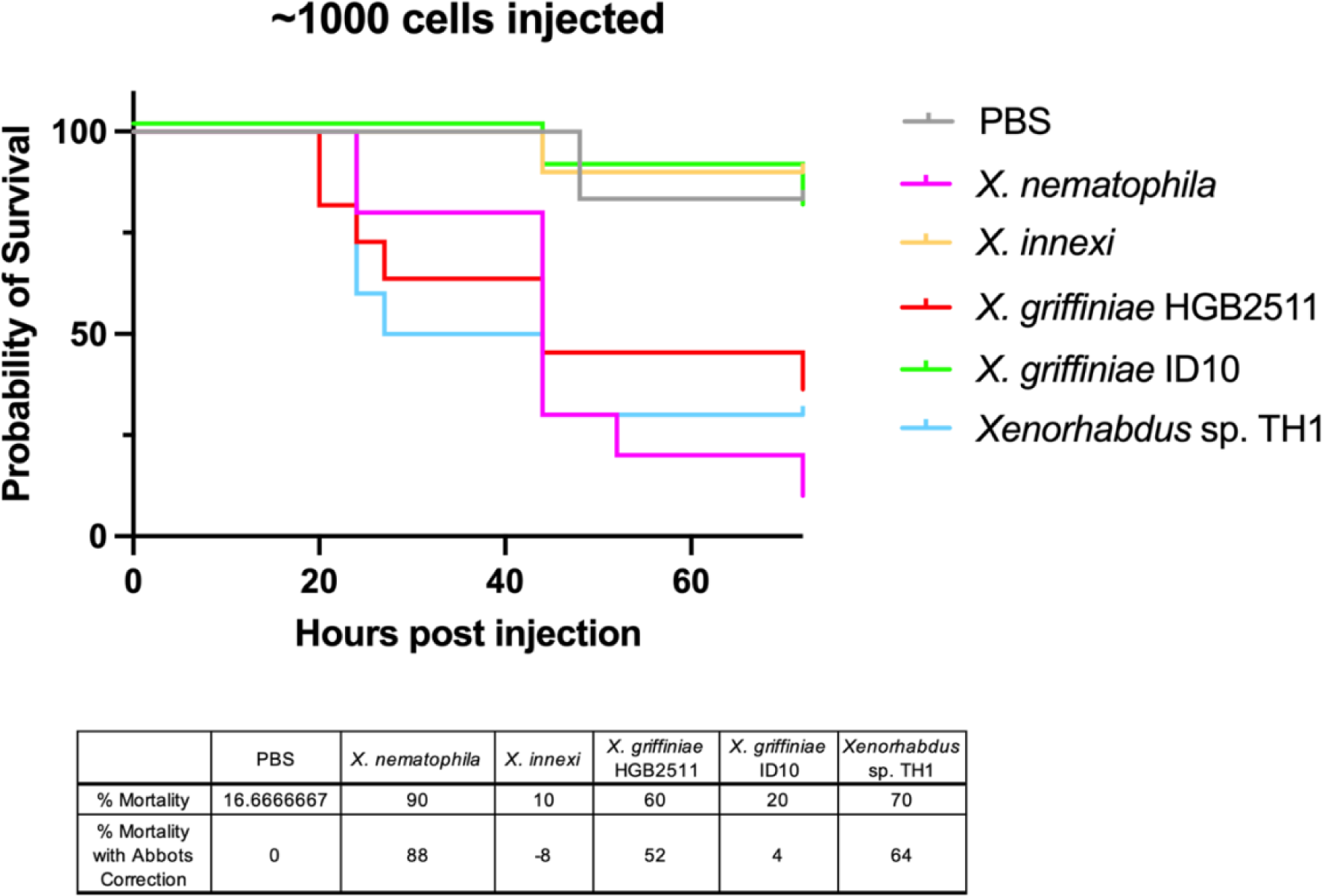
Comparative entomopathogenicity of *Xenorhabdus* bacteria to *Manduca sexta* larvae. Lines represent survival curves for insect larvae injected with approximately 1000 cells of each strain. Values have been corrected with Abbot’s formula within the table below.

## Discussion

As part of their symbiotic and entomopathogenic lifecycle, all *Xenorhabdus* must colonize and be transported by a nematode host, suppress insect immunity, establish a community within and consume the cadaver, and support reproduction of the nematode to ensure future transport [10]. Many will also compete or cooperate with other resident or transient microbial community members and respond to variations in abiotic factors and higher-order trophic interactions. Here we conducted a comparative genomics analysis to gain insights into the consequences of such variable selective pressures on the evolution of *Xenorhabdus* genome content. Our analysis centred around seven related *Xenorhabdus* strains, chosen based on their close phylogenetic proximity to *X. griffiniae*. Strains of this *Xenorhabdus* species are particularly relevant to the development of laboratory model systems used to study nematode-bacteria symbiosis because they are symbionts of *S. hermaphroditum,* currently the most genetically tractable steinernematid [24, 120, 25].

To allow detailed comparative genome analyses, we assembled new high-quality, circularised genomes for three strains: *X. griffiniae* ID10**^T^**, HGB2511, and TH1 (Table 1) [27, 24, 77]. High-quality genomes were already available for four other strains previously identified as *X. griffiniae*: xg97, Kalro, XN45 and VH1 [16, 80]. HGB2511 and the Kenyan isolates were verified as *X. griffiniae* species, as defined by a genome with pairwise values for dDDH and ANI that are >70% [121] and >95% [48] thresholds, respectively, with that of the type strain ID10 (Figure 1) [27]. Consistent with previous findings [16], our analysis confirmed that strain BMMCB is not an *X. griffiniae* strain as originally designated [122]. Instead, we found that strain BMMCB is conspecific with *Xenorhabdus* sp. SF857 (Sheet S1 in Additional File 2), the recently described type strain of a novel species *X. bakwenae* [8]. Strain TH1 is not conspecific to ID10 nor any other type strain, making it a novel species within the *Xenorhabdus* genus. These conclusions were further supported by phylogenetic reconstructions which revealed that *Xenorhabdus* sp. TH1 does not cluster with *X. griffiniae* but shares a last common ancestor with the progenitor of the *X. griffiniae* clade (Fig. 1). From our findings on the evolution of gene content, we speculate that TH1 diverged, primarily through gene losses, from the progenitor of the *X. griffiniae* clade to ultimately form its species (Fig. 5), similar to the speciation observed in *Bordetella pertussis* [123].

Phage-related genes are known drivers of genome variation between closely related strains [124] and previously were implicated as drivers of the differences in gene content among other close *Xenorhabdus* relatives of *X. griffiniae* [16]. The *X. griffiniae* genomes we analysed were found to be enriched in mobilome content, with phage-related genes specifically driving the diversification of subspecies and strains. We found that the India-Indonesia and Kenyan subspecies arose from the net gain of genes, of which 39-47% were from prophages (Fig. 4&5). Further, we found that the proportion of *X. griffiniae* strain-specific genes of prophage origin is 35-48%. This is especially high, considering that for example, of 30 *Bifidobacterium* strains examined, the highest proportion of strain-specific genes of prophage origin observed was only between 0.03-35.4% [125]. Moreover, *X. griffiniae* prophage regions reduced pairwise dDDH values among them by three percentage points. In *Salmonella enterica* prophage sequences have been shown to be highly variable and differentially conserved among strains, making them key drivers of genome diversification and useful markers for serovar typing [126, 127], Similarly, we conclude that in *X. griffiniae* prophages are major drivers of subspeciation and strain differentiation.

An essential component of the *Xenorhabdus* lifestyle is interaction with other organisms, including competing microbes, the mutualistic nematode host, and the prey insect host. Among the molecular machines that facilitate bacterial manipulation of other organisms is the T6SS, which delivers effectors into target (non-self) cells [128]. In this study, we found a high sequence similarity of corresponding *X. griffiniae* T6SS-encoding loci, but gene content variability in loci encoding concomitant effector proteins. This indicates that effector proteins possibly contribute to traits that vary within a species. One of these traits is nematode host specificity, which varies within both *X. griffiniae* and *X. bovienii* species [7]. In *X. bovienii*, the inactivation of *vgrG* in XG2-T6SS loci in strain SS-2004 resulted in the near loss (200-fold decrease) of the bacterium’s capacity to colonise its nematode host [129]. Although we found that the presence and sequence of VgrG does not appear to vary across *X. griffiniae* (or other *Xenorhabdus*), the genes encoding the effector proteins transported by VgrG do vary among the *X. griffiniae* species. Therefore, we speculate that one or more *X. griffiniae* VgrG-associated effector proteins may determine a strain’s capacity to naturally colonise its specific nematode host, similar to the conclusion reached about the T6SS function in *X. bovienii* SS-2004 [129].

CRISPR content and spacer identities support the conclusion that the strains studied here are diversifying due to phage pressure and reflect the taxonomic relationships we observed. The Kenyan subclade has nearly identical CIRSPR-Cas loci, consistent with their very close relationship. The only difference is that the XN45 CRISPR array 1a appears to lack a repeated spacer that the others have. Considering the Kenyan subclade as a single group, all the genomes were distinct from each other with respect to CRISPR spacer content, indicative of their unique histories in exposure to, and successful defence from phages and mobile genetic elements. Consistent with this, non-self-targeting protospacers identified within the group could be found within prophage genes and conjugation machinery in the genomes of the other strains. Our analysis identified instances of potential self-targeting, which offered the opportunity to search for anti-CRISPR genes, which are a key aspect of the co-evolution of phage and defence systems and have potential utility in applications of CRISPR technologies [94]. In the Kenyan subclade, we identified one clear candidate for such an anti-CRISPR locus comprising an HTH-domain Aca regulatory candidate and a small DUF2247 domain-containing protein ORF of 171 amino acids (aa), adjacent to the protospacer-containing gene *palA/fhaB,* which is also in the same region as a CRISPR array. However, DUF2247 proteins are also known as “imm38” and their presence within polymorphic toxin loci, such as *palA/fhaB,* has implicated them as immunity proteins to these toxins [130]. Consistent with this possibility, members of the Kenyan clade are the only strains of those analysed here that appear to have both a DUF2247 ORF and full-length *palA/fhaB* genes in that locus.

The *X. griffiniae* and *Xenorhabdus* sp. TH1 genomes we compared all bear hallmarks of the entomopathogenicity characteristic of the *Xenorhabdus* genus. However, when cultures were injected into *M. sexta* insect larva, the ID10 strain displayed attenuated virulence when compared with HGB2511 and TH1. The magnitude of this difference was directly comparable to the difference in virulence observed between *X. innexi* and *X. nematophila* species [68]. Members of the *X. bovienii* species group have demonstrated a similar range of virulence phenotypes when injected in the absence of the vectoring nematodes [131]. These differences may be due to genomic variation between the closely related species. Notably, we identified strain specific toxin loci, such as the *hcnABC* locus in TH1 and the zot domain containing toxins in the Kenyan clade that may underly different mechanisms or levels of virulence in *X. griffiniae* and its closest relative (Fig. 10; Table 3). It is also possible that the lack of virulence observed for ID10 is due to gene expression programs which control phenotypic variation locking the isolate in a state of attenuated virulence [132, 133]. If so, we predict that other isolates of the ID10 strain may retain high levels of virulence, similar to what has been observed for other *Xenorhabdus* species [132, 134, 135]. Alternatively, the ecological insect host range of ID10 may be distinctive enough from other *X. griffiniae* that it has lost the ability to infect the Lepidopteran insect *Manduca sexta* that we tested here.

## Conclusion

This study yielded three complete genome assemblies, which were of *X. griffiniae* ID10, *X. griffiniae* HGB2511 and *Xenorhabdus* sp. TH1. *Xenorhabdus* sp. TH1 is a novel bacterial species while *X. griffiniae* contained two subspecies. Both CRISPR loci and loci encoding T6SS effector proteins divided along these *X. griffiniae* subspecies lines. Intraspecific variation, including subspeciation, was largely driven by prophages. In terms of biosynthetic potential, *X. griffiniae* genomes encoded the production pathways of diverse and biotechnologically useful natural products such as antibacterials, antiprotozoals, and insecticidal toxins. Intraspecific variation in biosynthetic potential was observed, which we substantiated by the different levels of entomopathogenicity, among *X. griffiniae* strains, to *M. sexta*. Ultimately, these genome assemblies and genomic insights are foundational for continuing studies into the symbiosis between *X. griffiniae* and its self-fertilizing nematode host, *S. hermaphroditum*.

## Availability of data and materials

The datasets generated during the current study are available in the GenBank repository BioProject ID: PRJNA1085699. Previously reported data is available from public repositories (see Methods), and additional data is provided within the manuscript or supplementary information files.

## Authors’ contributions

JKH, RMA, and HGB collected, and analysed the data, and wrote the original draft of the manuscript. MC and GC performed DNA extraction, sequencing, and assembly for HGB2511 and TH1 genomes. JKH performed all other “bench” experiments, extractions and assemblies, with help from JM on the insect virulence assay. HGB supervised and provided resources for the study. All authors reviewed the manuscript.

## Acknowledgements

The authors would like to thank and acknowledge Zachary Burcham for assistance running the PathoFact software used in the toxin analysis and Sarah Kauffman for assistance with insect injections.

