## Additional File 1 for "Analyses of *Xenorhabdus griffiniae* genomes reveal two distinct sub-species that display intra-species variation due to prophages"

### Additional file 1: Additional results from analyses of secretion systems, CRISPR loci, maximum likelihood phylogenies, prophages and loci encoding type six effector proteins

**Table S1:** Comparison of the number of putative secretion systems that are present *Xenorhabdus griffiniae* strains

| | ***Xenorhabdus griffiniae*** | | | | | | ***Xenorhabdus* sp. TH1** | ***X. nematophila* 19061** |
|---|---|---|---|---|---|---|---|---|
| **Secretion System** | **HGB2511** | **ID10** | **Kalro** | **VH1** | **XN45** | **Xg97** | | |
| Flagellum | 1 | 1 | 1 | 1 | 1 | 1 | 1 | 1 |
| T1SS | 3 | 7 | 5 | 4 | 5 | 5 | 4 | 4 |
| T4aP | 1 | 1 | 1 | 1 | 1 | 1 | 1 | 1 |
| T5aSS | 2 | 2 | 2 | 2 | 2 | 2 | 2 | 1 |
| T5bSS | 1 | 1 | 1 | 1 | 1 | 1 | 1 | 1 |
| T6SS | 2 | 2 | 2 | 2 | 2 | 2 | 0 | 2 |

Definition of abbreviations: T1SS, type one secretion system; T5SS, type five secretion system; T6SS, type six secretion system; Flagellum, flagellum secretion system; T4aP, Type 4a pilus

**Table S2.** Loci encoding putative restriction-modification systems identified by homology and genome annotation searches. Alternating grey and white boxes indicate loci that are predicted to act together (Type I, II, and III restriction enzymes which are known to require multiple loci).

| Genome | Locus tag | Description | Method of Identification |
|---|---|---|---|
| BMMCB | LDNM01_v1_270038 | Type II Adenine-specific methyltransferase activity Eco57IA | BLASTP |
| BMMCB | LDNM01_v1_270039 | Type II Modification methylase Eco57IB | BLASTP |
| BMMCB | LDNM01_v1_450003 | Type I restriction-modification system, restriction subunit R | MAGE+BLASTP |
| BMMCB | LDNM01_v1_450006 | Type I restriction-modification system, specificity subunit S | MAGE+BLASTP |
| BMMCB | LDNM01_v1_450007 | putative Site-specific DNA-methyltransferase (adenine-specific) | BLASTP |
| BMMCB | LDNM01_v1_760025 | Site-specific DNA-methyltransferase (adenine-specific) | BLASTP |
| BMMCB | LDNM01_v1_760026 | Type I restriction-modification system, specificity subunit S | MAGE+BLASTP |
| BMMCB | LDNM01_v1_760028 | Type I site-specific deoxyribonuclease | BLASTP |
| BMMCB | LDNM01_v1_1590002 | DNA methyltrasferase (cytosine) dcm | BLASTP |
| BMMCB | LDNM01_v1_1590003 | DNA mismatch endonuclease VSP mismatch repair pathway | PROXIMITY |
| BMMCB | LDNM01_v1_1590004 | Type II Restriction endonuclease | MAGE |
| HGB2511 | XGHIN1_v1_3075 | Antirestriction protein | MAGE |
| HGB2511 | XGHIN1_v1_3130 | Type II restriction endonuclease | MAGE |
| HGB2511 | XGHIN1_v1_3131 | DNA mismatch endonuclease Vsr | PROXIMITY |
| HGB2511 | XGHIN1_v1_3132 | DNA-cytosine methyltransferase | BLASTP |
| HGB2511 | XGHIN1_v1_4004 | Type III restriction enzyme | MAGE+BLASTP |
| HGB2511 | XGHIN1_v1_4005 | adenine-specific DNA-methyltransferase | BLASTP |
| HGB2511 | XGHIN1_v1_3979 | putative restriction endonuclease hnh | MAGE |
| ID10 | XGHID_v1_1049 | Antirestriction protein | MAGE |
| ID10 | XGHID_v1_1232 | Antirestriction protein | MAGE |
| ID10 | XGHID_v1_1233 | Antirestriction plasmid protein | MAGE |
| ID10 | XGHID_v1_1327 | Antirestriction protein | MAGE |
| ID10 | XGHID_v1_1328 | Antirestriction protein | MAGE |
| ID10 | XGHID_v1_1421 | Antirestriction protein | MAGE |
| ID10 | XGHID_v1_1422 | Antirestriction protein | MAGE |
| ID10 | XGHID_v1_0644 | Type I restriction enzyme Methylase protein | MAGE+BLASTP |
| ID10 | XGHID_v1_0645 | Type I restriction enzyme, S subunit | MAGE+BLASTP |
| ID10 | XGHID_v1_0646 | Type I restriction enzyme, R subunit | MAGE+BLASTP |
| ID10 | XGHID_v1_1278 | Type II restriction endonuclease | MAGE |
| ID10 | XGHID_v1_1279 | DNA mismatch endonuclease Vsr | PROXIMITY |
| ID10 | XGHID_v1_1280 | DNA-cytosine methyltransferase dcm | BLASTP |
| ID10 | XGHID_v1_1102 | Type IV restriction system protein | MAGE |
| ID10 | XGHID_v1_3086 | predicted Type IV restriction endonuclease | MAGE |
| ID10 | XGHID_v1_4283 | putative restriction endonuclease hnh | MAGE |
| Kalro | JASDYB01_13172 | Antirestriction protein | MAGE |
| Kalro | JASDYB01_13173 | Antirestriction protein | MAGE |
| Kalro | JASDYB01_14510 | Type I restriction modification DNA specificity domain-containing protein | MAGE+BLASTP |
| Kalro | JASDYB01_14511 | Adenine-specific methyltransferase activity bcglA | BLASTP |
| Kalro | JASDYB01_10684 | Type IV Restriction endonuclease | MAGE |
| Kalro | JASDYB01_13223 | predicted Type IV restriction system protein | MAGE |
| TH1 | XTH1_v2_2577 | Antirestriction protein | MAGE |
| TH1 | XTH1_v2_0732 | Type I restriction enzyme EcoR124II Methylase protein hsdM | MAGE+BLASTP |
| TH1 | XTH1_v2_0733 | Type I restriction enzyme, S subunit | MAGE+BLASTP |
| TH1 | XTH1_v2_0735 | Type I restriction enzyme EcoR124II R protein hsdR | MAGE+BLASTP |
| TH1 | XTH1_v2_3085 | putative Type I restriction enzyme HindVIIP Methylase protein | MAGE+BLASTP |
| TH1 | XTH1_v2_3086 | Type I restriction enzyme, S subunit | MAGE+BLASTP |
| TH1 | XTH1_v2_3087 | Type I restriction enzyme, R subunit | MAGE+BLASTP |
| TH1 | XTH1_v2_1298 | predicted Type IV restriction endonuclease | MAGE |

**Table S3:** Biosynthetic gene clusters (BGCs) in the *Xenorhabdus griffiniae* ID10 genome

| BGC | Type | Biosynthesis it encodes | Putative Bioactivity | ID10 genome locus | HGB2511 genome locus |
|---|---|---|---|---|---|
| | | | Known BGCs | | |
| *safA-H* | NRPS | Safracin | Antibacterial | 119,533-137,302 | 129,186-118,464 |
| *lpcS* | NRPS | Group IIA Bovienimides | Insect immunity suppressor | 2,775,620-2,787,196 | 1,764,757 – 1,776,333 |
| *apeA-R* | PKS | Arylpolyenes | | 4,052,973-4,067,609 | 600,909 – 586,271 |
| *isnAB, GT* | Tyrosine derivative | Rhabduscin | Insect immunity suppressor | 3,276,469-3,279,680 | Not present |
| *xvbA-J* | NRPS | Benzobactin | Cytotoxicity | 1,343,066-1,325,877 | Not present |
| *ioc/leu* | ß-lactone | IOC | Cytotoxicity | 3,604,901-3,611,464 | 1,024,776-1,019,356 |
| *bcmA-G* | CDPS | Bicyclomycin | Antibacterial | 3,074,545-3,081,355 | Not present |
| *gxpS* | NRPS | GameXpeptide C | Insect immunity suppressor | 457,463-472,960 | 458,227–473,709 |
| *paxABC* | NRPS | Novel PAX peptides | Antifungal, antibacterial | 3,113,099-3,140,411 | 1,447,169 - 1,437,220 |
| *xabA-D* | NRPS | Novel xenoamicins | Antiprotozoal | 2,042,213-1,996,480 | 2,520,649-2,566,406 |
| *darA-E* | RIPP | Novel darobactin | antibacterial | 2,219,063-2,212,963 | 2,353,462-2,359,501 |
| *pxbA-J* | NRP-metallo-phore | Photo-xenobactin | Metallophore & insecticide | 2,463,657-2,494,495 | 2,104,858 – 2,074,029 |
| *tnbA-F* | NRP-metallo-phore | Triscatechol siderophores | Siderophore | 573,831-592,583 | 3,958,982-3,940,260 |
| *spe* | Amino-glycoside | Actino-spectacin | Antibacterial | 1,136,403-1,147,206 | Not present |

| | | | | | |
|---|---|---|---|---|---|
| | | | Unknown BGCs | | |
| | NRPS | *N*-terminally acylated unodo-decapeptide* | | 1,973,520-1,958,806 | 2,566,703-2,607,363 |
| | NRPS | Hse-His containing peptide | | 3,288,877-3,296,952 | Not present |
| | NRPS | *N*-terminally acylated $^{D}$Gln-$^{L}$Trp-$^{D}$Tyr-$^{D}$Tyr-$^{L}$Trp-$^{L}$Tyr | | Not present | 2,067,796-2,043,896 |
| | NRPS-PKS | Gln & Asp containing NRP-PKS hybrid | | 1,382,055-1,401,288 | Not present |
| | NRPS-PKS | Arg-Asp containing peptide polyketide hybrid | | 2,186,186-2,207,667 | Not present |
| | RIPP | Thiopeptide | | 1,790,808-1,784,605 | 2,855,135 -2,861,338 |
| | RIPP | Thioamitides | | 1,304,216-1,314,671 | 3,899,202-3,909,658 |
| | Phospho-nate | | | 623,023-613,019 | 3,923,655-3,913,553 |
| | ß-lactone | | | 2,663,491-2,655,349 | 1,826,388-1,832,145 |

NRPS, non-ribosomal peptide synthetase; PKS, polyketide synthase; RIPP, ribosomally synthesized and post-translationally modified peptides; CDPS, cyclic dipeptide synthase; IOC, 3-isopropyl-4-oxo-2-oxetanecarboxylic acid; PAX, peptide antimicrobials from *Xenorhabdus*. *For the HGB2511 strain, the homologous BGC is predicted to encode the production of a dodecapeptide derivative.

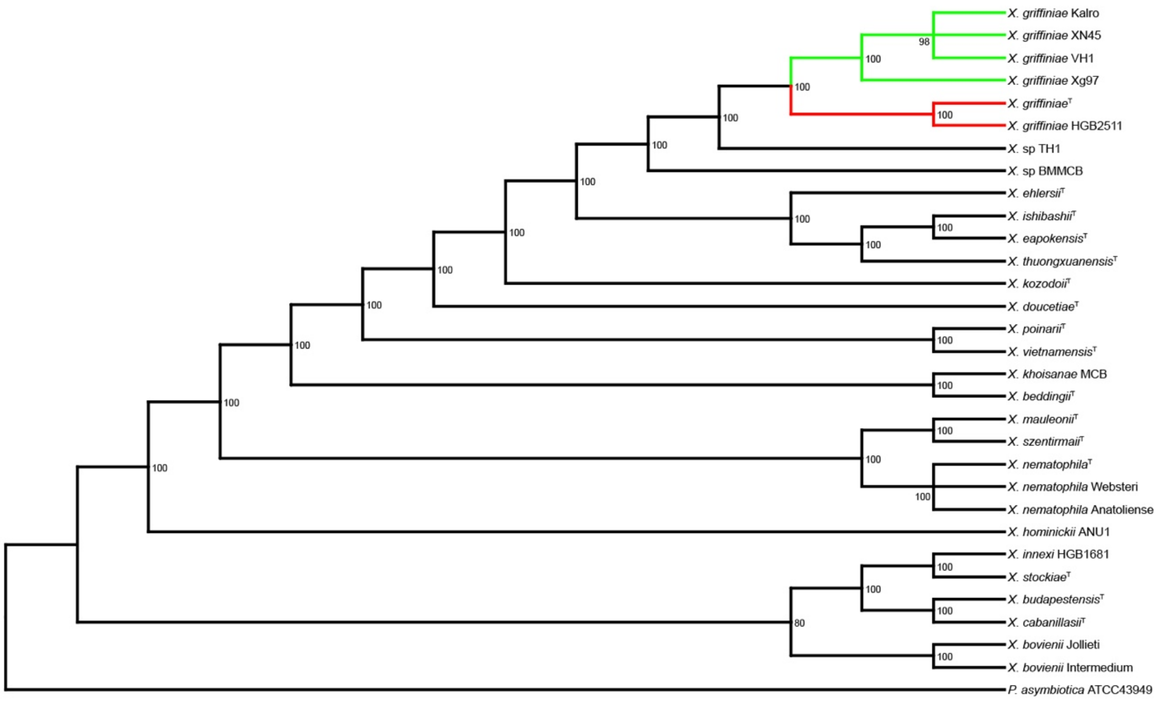


**Figure S1:** Maximum likelihood phylogenetic tree created using RAxML with one-to-one orthologs from *Xenorhabdus* type strains, closely related *X. griffiniae* strains, and *Photorhabdus asymbiotica* as an outgroup. Bootstrapping was performed with 1,000 replicates and nodes with support less than 75 were collapsed. Strains of *X. griffiniae* formed two separate clades, those isolated from *S. hermaphroditum* nematodes found in Indonesia (ID10) and India (HGB2511) (red) and those isolated from nematodes found in Kenya (green).

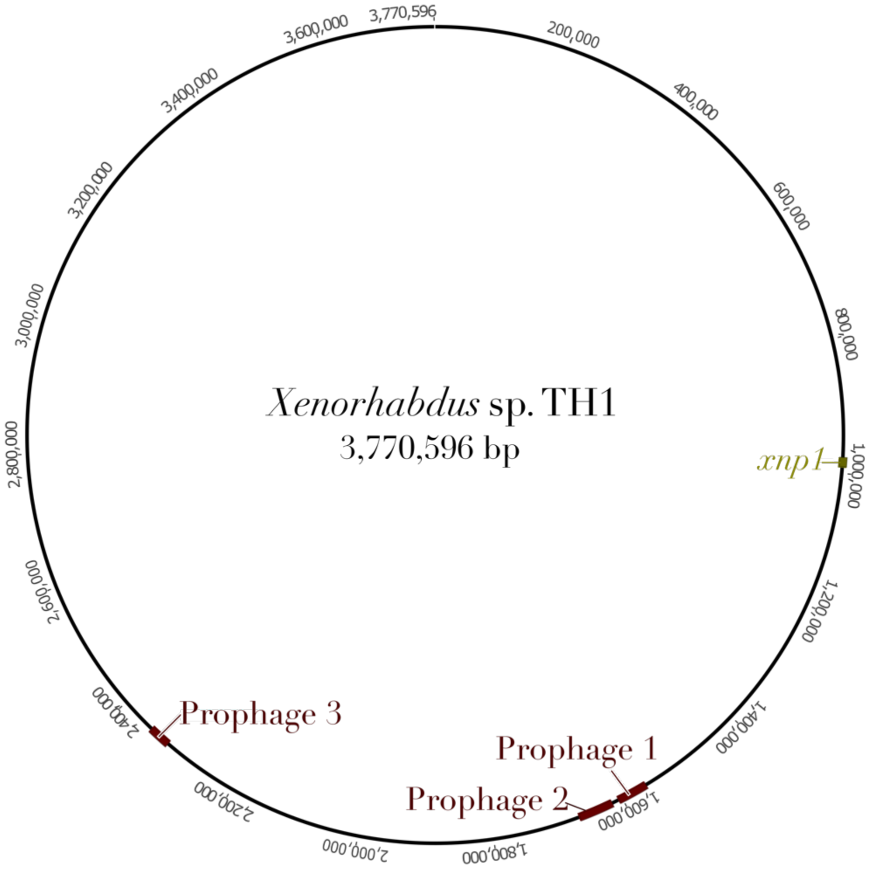


**Figure S2**: Loci of prophages and *xnp1* in the complete *Xenorhabdus* sp. TH1 genome

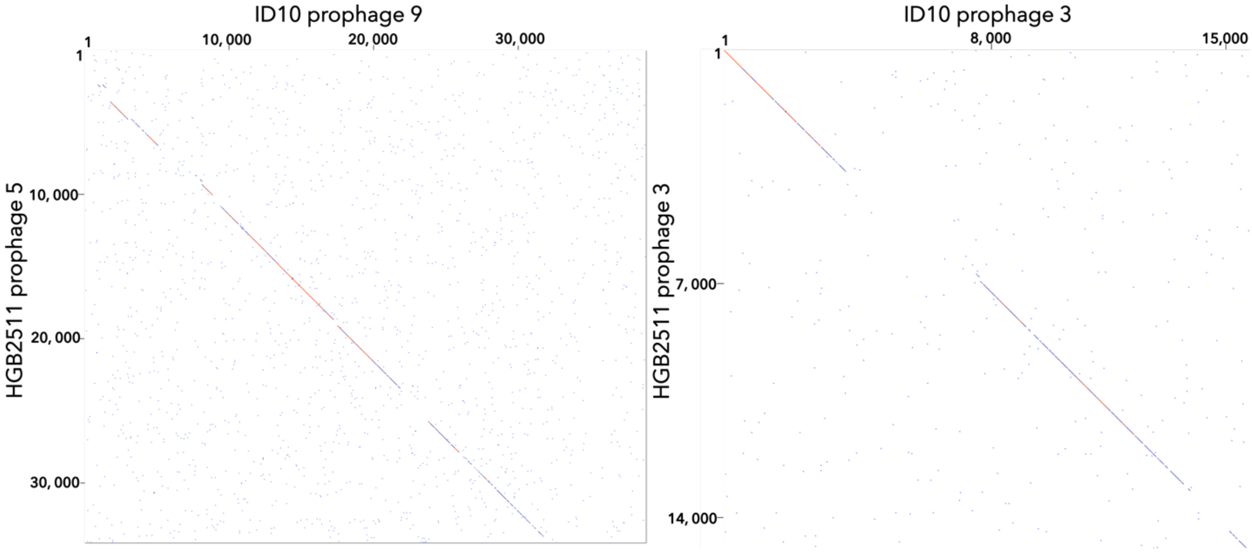


**Figure S3**: Dotplots of prophage loci that were considerably similar between strains.

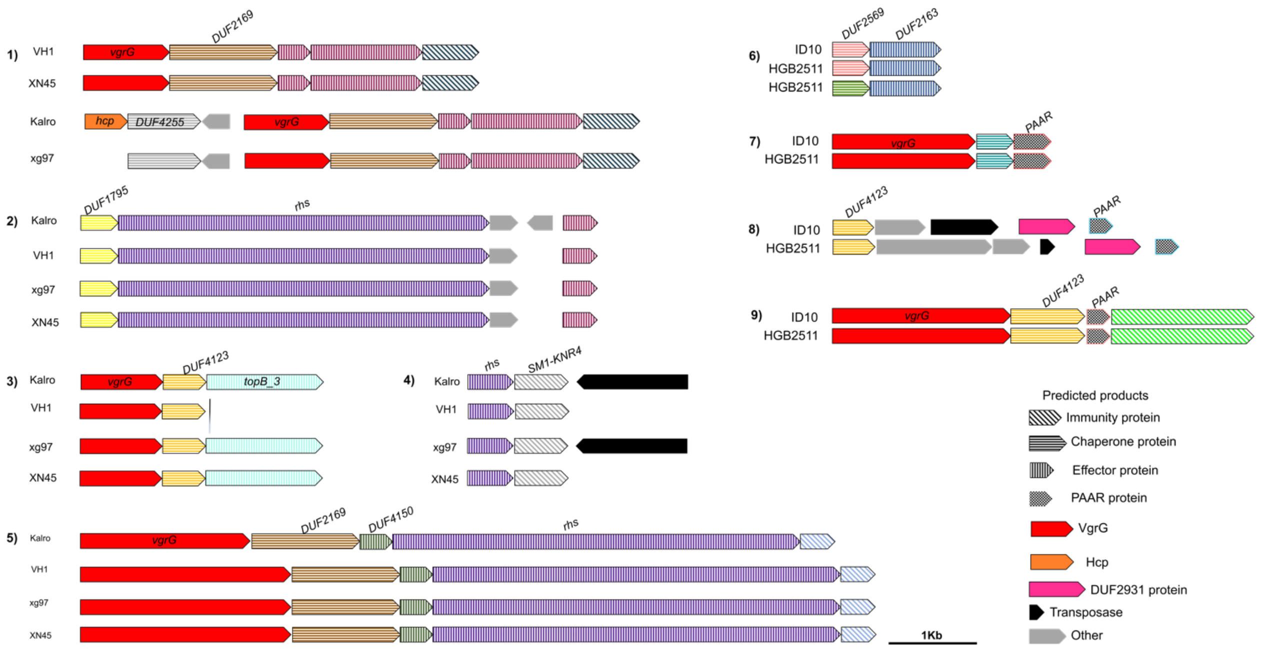


**Figure S4**: Schematic of subspecies-specific type six secretion system (T6SS) effector-encoding loci in six strains of *Xenorhabdus griffiniae*. Loci 1-5 are specific to the Kenyan subspecies while loci 6-9 are specific to the India-Indonesia subspecies.

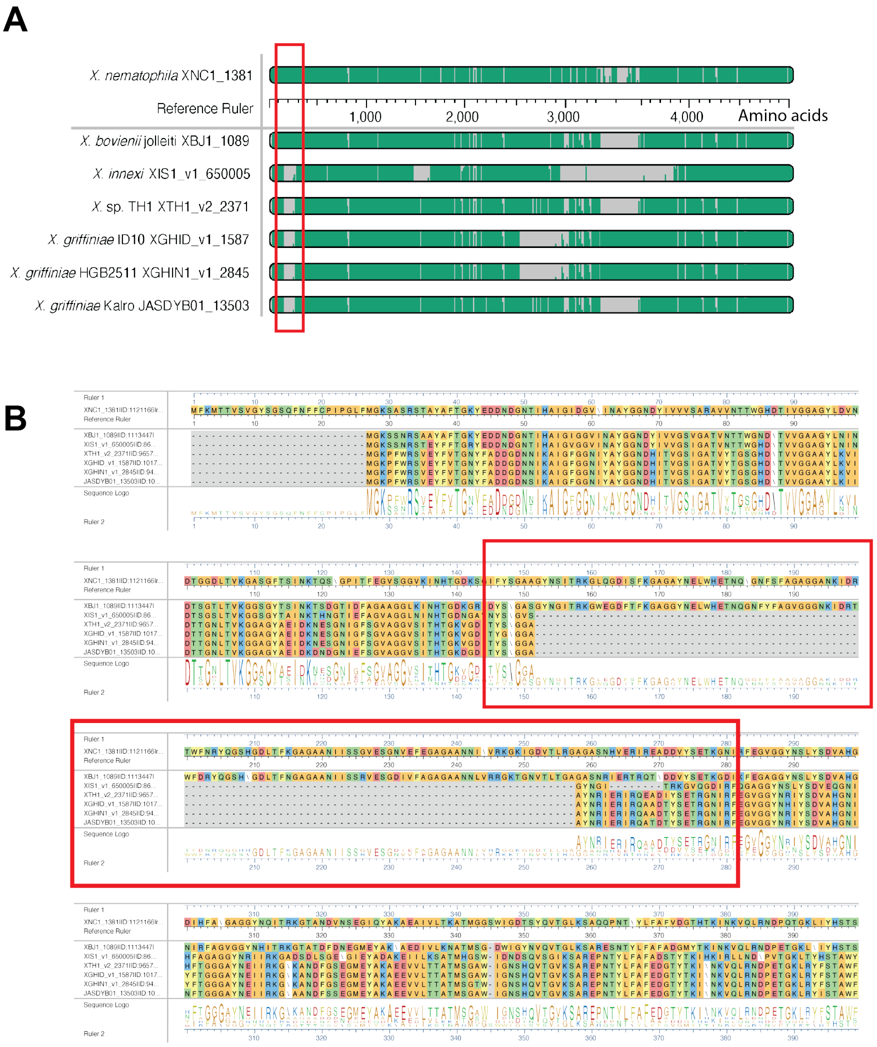


**Figure S5:** Overview of MARTX proteins from HGB2511, ID10, Kalro and TH1 (XGHIN1_v1_2845, XGHID_v1_1587, XTH1_v2_2371, JASDYB01_13503) and *X. innexi* XIS1_v1_650005 aligned to *X. nematophila* MARTX locus XNC1_1381 as reference using MUSCLE **A)** Overview of whole protein alignment. Grey bars indicate gaps in the sequence alignment and the red box shows the gap in A repeats at the N-terminus of the *X. innexi* protein sequence which is also observed in the *Xenorhabdus* sp. TH1, and *X griffiniae* sequences. **B)** Detailed amino acid alignment of the MARTX N-terminus (1-400aa) showing gaps in the region containing the A repeats present in the *X.* sp TH1, and *X griffiniae* sequences (red box: position ~150-200aa).

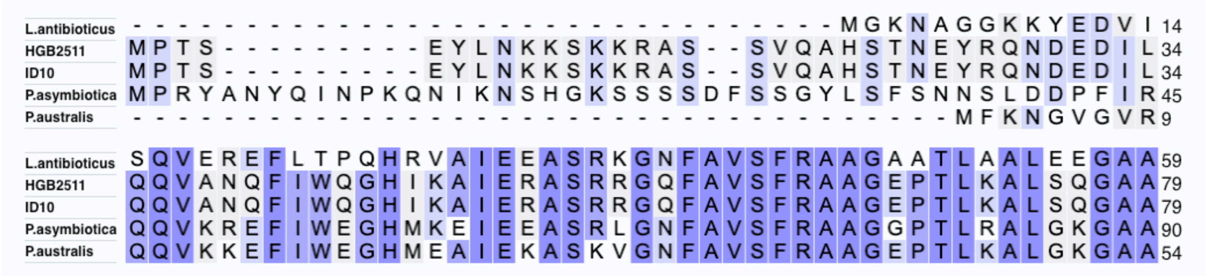


**Figure S6:** *N*-terminus of the Clustal Omega multiple sequence alignment of the Pdp1 homologs from ID10, HGB2511, *Photorhabdus asymbiotica, Photorhabdus australis, and Lysobacter antibioticus.*

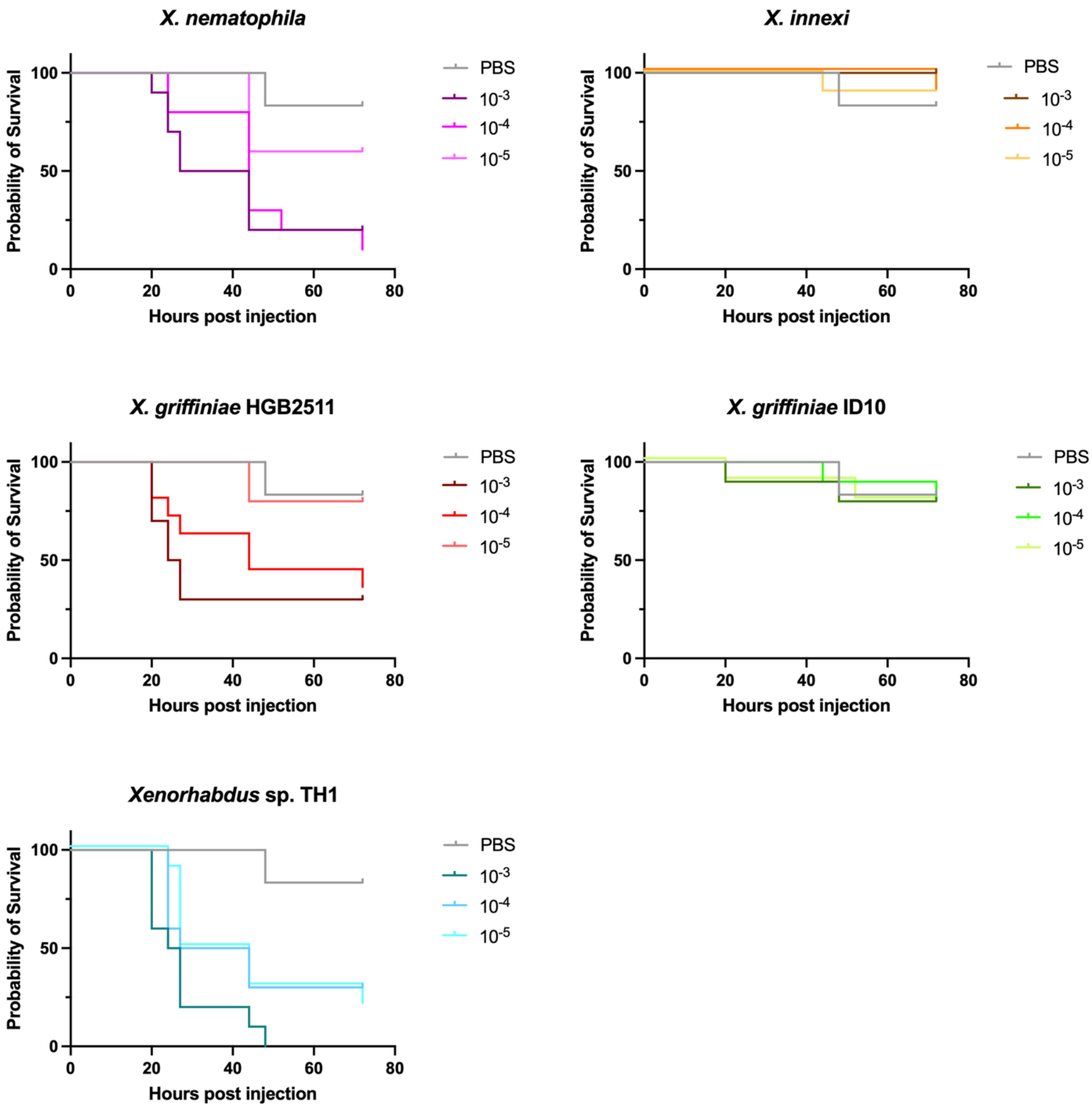


**Figure S6:** Percentage survival of *Manduca sexta* larvae post injection with live *Xenorhabdus* bacteria cells. Lines represent survival curves for insect larvae injected with 10 μl of PBS and $10^{-3}$, $10^{-4}$, and $10^{-5}$ PBS dilutions of log-phase bacterial cultures.
