## Additional File 3 for "Analyses of *Xenorhabdus griffiniae* genomes reveal two distinct sub-species that display intra-species variation due to prophages"

### Additional file 3. Word document with additional details of the detection and analysis of *Xenorhabdus griffiniae* CRISPR loci

**Table S4**. Locus tags and coordinates of CRISPR-Cas features. The MaGe locus tags are listed for genes encoded at *cas* loci (region 1 for all genomes, and region 2 for HGB2511, ID10, and TH1) in each of the analyzed strains. Annotated gene names are provided on the left (for region 1) and on the right (for region 2) of the table. The genome start coordinate of the start codon for the *cas3* gene is provided at the bottom of the *cas* list. For each CRISPR repeat region found in a genome, the start coordinate of the first repeat and the number of spacers identified are provided.

| | | HGB2511 XGHIN1 | | ID10 Xg_ID | | xg97 JASDYA01 | Kalro JASDYB01 | XN45 JACWFC01 | VH1 JADEUF01 | TH1 XTH1 | | BMMCB LDNM01 | |
|---|---|---|---|---|---|---|---|---|---|---|---|---|---|
| | Region 1 annotation | Region 1 | Region 2 | Region 1 | Region 2 | Region 1 | Region 1 | Region 1 | Region 1 | Region 1 | Region 2 | Region 1 | Region 2 annotation |
| | bleo/gloA | XGHIN1_v1_0694 | XGHIN1_v1_3887 | XGHID_v1_3830 | XGHID_v1_4178 | JASDYA01_11181 | JASDYB01_11232 | JACWFC01_130039 | JADEUF01_2300036 | XTH1_v2_3015 | XTH1_v2_3073 | LDNM01_v1_310094 | eda |
| | cas3 | XGHIN1_v1_0693 | XGHIN1_v1_3886 | XGHID_v1_3831 | XGHID_v1_4177 | JASDYA01_11182 | JASDYB01_11233 | JACWFC01_130040 | JADEUF01_2300037 | XTH1_v2_3016 | XTH1_v2_3072 | LDNM01_v1_310093 | cas3 |
| | casA/cse1 | XGHIN1_v1_0692 | XGHIN1_v1_3884 | XGHID_v1_3832 | XGHID_v1_4175 | JASDYA01_11183 | JASDYB01_11234 | JACWFC01_130041 | JADEUF01_2300038 | XTH1_v2_3017 | XTH1_v2_3070 | LDNM01_v1_310092 | casA/cse1 |
| | casB/cse2 | XGHIN1_v1_0691 | XGHIN1_v1_3883 | XGHID_v1_3833 | XGHID_v1_4174 | JASDYA01_11184 | JASDYB01_11235 | JACWFC01_130042 | JADEUF01_2300039 | XTH1_v2_3018 | XTH1_v2_3069 | LDNM01_v1_310091 | casB/cse2 |
| | casC/cse4 | XGHIN1_v1_0690 | XGHIN1_v1_3882 | XGHID_v1_3834 | XGHID_v1_4173 | JASDYA01_11185 | JASDYB01_11236 | JACWFC01_130043 | JADEUF01_2300040 | XTH1_v2_3019 | XTH1_v2_3068 | LDNM01_v1_310090 | casC/cse4 |
| | casD | XGHIN1_v1_0689 | XGHIN1_v1_3881 | XGHID_v1_3835 | XGHID_v1_4172 | JASDYA01_11186 | JASDYB01_11237 | JACWFC01_130044 | JADEUF01_2300041 | XTH1_v2_3020 | XTH1_v2_3067 | XXXXX | casD |
| | casE/cas5 | XGHIN1_v1_0688 | XGHIN1_v1_3880 | XGHID_v1_3836 | XGHID_v1_4171 | JASDYA01_11187 | JASDYB01_11238 | JACWFC01_130045 | JADEUF01_2300042 | XTH1_v2_3021 | XTH1_v2_3066 | LDNM01_v1_310088 | casE/cas5 |
| | cas1/cse3 | XGHIN1_v1_0687 | | XGHID_v1_3837 | | JASDYA01_11188 | JASDYB01_11239 | JACWFC01_130046 | JADEUF01_2300043 | XTH1_v2_3022 | | LDNM01_v1_310087 | |
| | cas2 | XGHIN1_v1_0686 | | XGHID_v1_3838 | | JASDYA01_11189 | JASDYB01_11240 | JACWFC01_130047 | JADEUF01_2300044 | XTH1_v2_3023 | | XXXXX | |
| | lipoprotein | XGHIN1_v1_0685 | | XGHID_v1_3839 | | JASDYA01_11190 | JASDYB01_11241 | JACWFC01_130048 | JADEUF01_2300045 | XTH1_v2_3024 | | LDNM01_v1_310085 | |
| | membrane protein | XGHIN1_v1_0684 | | XGHID_v1_3840 | | JASDYA01_11191 | JASDYB01_11242 | JACWFC01_130049 | JADEUF01_2300046 | XTH1_v2_3025 | | LDNM01_v1_310084 | |
| cas3 | start coordinate | 687309 | 4082009 | 3918399 | 4212171 | 1149686 | 1175147 | 358704 | 3279755 | 3277536 | 3342027 | 1956764 | |
| Repeat Region 1a | start coordinate | 689662 | | 3917418 | | 1147059 | 1172520 | 356138 | 3277128 | 3275744 | | 1958872 | |
| | Spacer # | 23 | | 1 | | 28 | 28 | 28 | 28 | 31 | | 6 | |
| Repeat Region 1aii | start coordinate | 678652 | | 3927056 | | 1158349 | 1183810 | 367367 | 3288418 | 3927056 | | NA | |
| | Spacer # | 0 | | 0 | | 0 | 0 | 0 | 0 | 0 | | NA | |
| Repeat region 1bi | start coordinate | 4079238 | | 4209400 | | NA | NA | NA | NA | 3339253 | | NA | |
| | Spacer # | 0 | | 0 | | NA | NA | NA | NA | 0 | | NA | |
| Repeat region 1bii | start coordinate | 4074160 | | 4204292 | | 2327296 | 2359571 | 3473979 | 3529539 | 3334252 | | NA | |
| | Spacer # | 24 | | 1 | | 8 | 8 | 8 | 8 | 35 | | NA | |
| Repeat region 2 | start coordinate | 2136087 | | 2440560 | | 4207989 | 4246909 | 790425 | 2634716 | 2090802 | | NA | |
| | Spacer # | 13 | | 0 | | 18 | 18 | 18 | 18 | 51 | | NA | |

**Table S5**. Representative putative protospacers of *Xenorhabdus* strain CRISPR spacers, identified using sequence similarity searches in the CRISPRTarget platform (See Materials and Methods in main text). Shown are source strain of CRISPR spacer (Strain) and CRISPR spacer label (Spacer; See Fig. 8D-F and Table S4), and the annotation (Protospacer ID) and general category of annotated function (Protospacer Type) of each protospacer-containing locus. The blue highlighting indicates those spacers that also had protospacers among the closest relatives of *X. griffiniae*. The gold, green, and gray highlighting indicates protospacers from eukaryotic viruses, plasmids, or genes of unknown function, respectively. All other representative protospacers shown were in putative bacterial prophage genes.

| Strain | Spacer | Protospacer ID | Protospacer Type |
|---|---|---|---|
| HGB3511 | 1b-2 | MH673672.2_Thermus phage phiFa, complete genome. | Phage |
| | 1b-5 | Wenling_crustacean virus 10 strain WLJQ101844 hypothetical protein 1 | Crustacean virus |
| | 1b-7 | MH884508.1_Bacillus phage vB_BcoS-136, complete genome. | Phage |
| | 1b-9 | MT835525.1_UNVERIFIED: Bacteriophage sp. clone scaffold_96 genomic sequence, sequence. | Phage |
| | 1b-11 | Enterobacteria_phage Fels-2, complete genome | Phage |
| | 1b-16 | MZ188976.1_Cronobacter phage JK004, complete genome. | Phage |
| | 1b-17 | Streptomyces_sp. BHT-5-2 plasmid p1, complete sequence | Plasmid |
| | 1b-18 | Serratia_entomophila plasmid pADAP, complete sequence | Plasmid |
| | 1b-24 | OP172789.1_Escherichia phage ECO71P1, complete genome. | Phage |
| | 2-4 | KF021268.1_Staphylococcus phage philBB-SEP1, complete genome. | Phage |
| | 2-5 | KM289195.1_Streptococcus phage T12, complete genome | Phage |
| | 2-9 | Photobacterium_damselae subsp. piscicida plasmid pP99-018, complete sequence | Plasmid |
| | 2-10 | Methermicoccus_shengliensis DSM 18856 BP07DRAFT_scaffold00001.1_C, whole genome shotgun sequence | unknown |
| Kalro | 1a-1 | Gordonia_phage Maridalia, complete genome | Phage |
| | 1a-2 | FJ429185.1_Lactococcus phage P087, complete genome | Phage |
| | 1a-10 | CP025900.1_Escherichia phage 500465-1, complete genome | Phage |
| | 1a-19 | LN889991.1_Bacteriophage 15G genome assembly, chromosome: 1. | Phage |
| | 1a-22 | OX241423.1_Vibrio phage 249E41-1 genome assembly, chromosome: VP115E34_1. | Phage |
| | 1a-23 | Thiomonas_intermedia strain ATCC 15466 plasmid pXI1, complete sequence | Plasmid |
| | 1b-2 | Methanobrevibacter_boviskoreani JH1, whole genome shotgun sequence | Unknown |
| | 1b-4 | LC553736.1_Salmonella phage SAP012 DNA, complete sequence. | Phage |
| | 1b-7 | MN019128.1_Escherichia phage Henu7, complete genome. | Phage |
| | 1a-1 | CP069347.1_Clostridioides phage pCD5401_3, complete sequence. | Phage |
| TH1 | 1a-2 | AF063097.1_Bacteriophage P2, complete genome. | Phage |
| | 1a-10 | KR136260.1_Polaribacter phage P12002S, complete genome. | Phage |
| | 1a-26 | HM770079.1_Salmonella phage RE-2010, complete genome. | Phage |
| | 1a-29 | AP014629.1_Edwardsiella phage GF-2 DNA, complete sequence. | Phage |
| | 1a-31 | KM983332.1_Clostridium phage phiCT19406C, complete genome. | Phage |
| | 1b-17 | MG592612.1_Vibrio phage 1.247.A._10N.261.54.E12, partial genome. | Phage |
| | 1b-22 | MT836127.1_UNVERIFIED: Bacteriophage sp. clone scaffold_1474 genomic sequence, sequence. | Phage |
| | 1b-23 | KU686207.1_Synechococcus phage S-CAM22 isolate 0210CC35, complete genome. | Phage |
| | 2-2 | ON529857.1_Brevundimonas phage vB_BpoS-Kikimora, complete genome | Phage |
| | 2-5 | KY271401.1_Klebsiella phage 1 LV-2017, complete genome. | Phage |
| | 2-15 | OQ718158.1_Escherichia phage vB_Ec-M-J, complete genome | Phage |
| | 2-17 | Equid_herpesvirus 4, complete genome | Equid Virus |
| | 2-19 | AY135486.1_Salmonella phage PSP3, complete genome. | Phage |
| | 2-20 | CP000711.1_Enterobacteria phage CUS-3, complete genome | Phage |
| | 2-21 | OK138555.1_Klebsiella phage vB_Kp_IME328, complete genome. | Phage |
| | 2-24 | MK416011.1_Klebsiella phage ST437-OXA245phi4.1, complete genome. | Phage |
| | 2-25 | MK448918.1_Streptococcus phage Javan366, complete genome. | Phage |
| | 2-26 | proph_161422_GID_2878435 | Phage |
| | 2-31 | OM284015.1_Elizabethkingia phage EKP1, complete genome. | Phage |
| | 2-41 | MH051918.1_Escherichia phage vB_EcoS_IME347, complete genome. | Phage |
| | 2-43 | proph_160513_GID_2803451 | Phage |
| | 2-44 | OM835952.1_Klebsiella phage KP12 clone KP12_2, partial sequence. | Phage |
| | 2-47 | DQ426904.1_Mannheimia phage PHL101, complete genome. | Phage |
| | 2-48 | ON287372.1_Serratia phage vB_SmaS-Totoro, complete genome. | Phage |
| BMMCB | 1a-1 | Enterobacter_cloacae strain 20ES map unlocalized plasmid p20ES-132 tig00000049, whole genome shotgun sequence | Plasmid |

**Additional Information on CRISPR array identification in *X. griffiniae* and related strains**

HGB2511, Kalro, ID10, TH1, and BMMCB genomes were submitted to CRISPRdetect (http://crispr.otago.ac.nz/CRISPRDetect/ CRISPRDetect 2.3) to identify potential CRISPR regions. The program did not detect any repeats in ID10. The consensus repeat sequences (29 bp) for each of the identified regions (here referred to as Region 1a, Region 1b, and Region 2 – see main text) of the other strains is shown below. Variability within the region among strains is highlighted in green.

| | |
|---|---|
| Region 1a | |
| HGB2511 -1a | GTGTTCCCCGTAAGTACGGGGATAAACCG |
| Kalro -1a | GTGTTCCCCGTGAGTACGGGGATAAACCG |
| TH1 -1a | GTGTTCCCCGTGAGTACGGGGATAAACCG |
| BMMCB -1a | GTGTTCCCCGTGAGTACGGGGATAAACCG |
| Region 1b | |
| HGB2511 -1b | CTGTTCCCCATGTGTATGGGGATAAACCG |
| Kalro- 1b | CTGTTCCCCATATGTATGGGGATGAACCG |
| TH1 -1b | CTGTTCCCCATGTGTATGGGGATAAACCG |
| Region 2 | |
| HGB2511 -2 | GTGTTCCCCGTGAGTACGGGGATAAACCG |
| Kalro -2 | GTGTTCCCCGTGAGTACGGGGATAAACCG |
| TH1 -2 | GTGTTCCCCGTGAGTACGGGGATAAACCG |
| BMMCB -2 | ATGTTCCCCGCGTACGCGGGGATAAACCG |

A manual search within the 9 selected genomes (HGB2511, ID10, xg97, Kalro, XN45, VH1, and TH1) for CRISPR repeats was conducted. Blastn (query coverage >0% and identity >0%) was used to search for sequence similarity to the previously published *Xenorhabdus* consensus CRISPR repeat [1]. This search yielded the same results as CRISPRdetect with two exceptions: the manual search did not detect the BMMCB region 2 repeats found by CRISPRdetect, but it did reveal five potential CRISPR repeat regions in ID10 that were not found by CRISPRdetect. Two of these were in region 1a and are denoted 1ai and 1aii, two were in region 1b, denoted 1bi and 1bii, and one was in region 2. Each is predicted to encode at most one spacer. To further analyze the ID10 repeat regions, the sequences were aligned manually with the CRISPRdetect consensus repeat sequence from each region, shown below highlighted in blue, with a 32-bp spacer region indicated by “X”. In addition, the last repeat-spacer-repeat sequence of each region in HGB2511 was included for comparison, since these were called by CRISPRdetect. Differences with the consensus are noted with yellow highlighting.

```
Consensus      GTGTTCCCCGTGAGTACGGGGATAAACCGXXXXXXXXXXXXXXXXXXXXXXXXXXXXXXXXGTGTTCCCCGTGAGTACGGGGATAAACCG
HGB2511 -1a    GTGTTCCCCGTAAGTACGGGGATAAACCGCCCAATTCAGGCATAGTAAGACCCCGATAAACGTGTTCCCCGTAAGTACGGGGATAAACAC – Y
ID10 -1ai      GTGTTCCCCGTAAGTACGGGGATAAACCGGTAGTCAACCTCGGATGACAATTTACTTCTATGTGTTCCCCGTGAATACGGGGATAAACAC – Y
ID10 -1aii     GTGTTCCCTGTAGGTACGGGGATAAACCGCAAACGCTGATTATAGAACGAGACAAAAACGAATATTCCCCATAGGCTCAATGAATAATTC – N

Consensus      CTGTTCCCCATGTGTATGGGGATAAACCGXXXXXXXXXXXXXXXXXXXXXXXXXXXXXXXXCTGTTCCCCATGTGTATGGGGATAAACCG
HGB2511 -1b    CTGTTCCCCATGTGTATGGGGATAAACCGAATGGCGCGGTGGTTACTGTCGTGTATACATTCTGTTCCCCATGTGTATGGGGATAAACCG – Y
ID10 -1bi      GTATTCCCCGTGAGTACGGGGATAAACCGTCATTCTGTACATTAATAATGATGTTTATCATCAAGTTCCCAAACCCTTTTGGGATTTATT – N
ID10 -1bii     CTGTTCCCCATGTGTATGGGGATAAACCGGGTGCTACCTGAAATACGGGGTGGCAAATACCCTGTTCCCCATGTGTATGGGGATAAATCG – Y

Consensus      GTGTTCCCCGTGAGTACGGGGATAAACCGXXXXXXXXXXXXXXXXXXXXXXXXXXXXXXXXGTGTTCCCCGTGAGTACGGGGATAAACCG
HGB2511 -2     GTGTTCCCCGTGAGTACGGGGATAAACCGTATTTGTAGTTAGATCAGGCTGGACTCAGGCAGTGTTCCCCGTGAGTACGGGGATAAACCG – Y
ID10 -2        GTGTTCCCCGTGAGTACGGGGATAAACCGCGCATGAGCAGTTTAATTGAATATTCACAGGCGTGTTCCCTGTGAGTGTAGTGATGTGATG — N
```

Since ID10 1ai and ID10 1bii regions have the same or fewer differences from consensus as the last HGB2511 repeat for each region, we have included these as bona fide, single spacer CRISPR loci, with the spacer sequence highlighted in red. Consistent with this call, these CRISPR repeats and spacer are syntenic with the longer CRISPR arrays found in HGB2511, the Kenyan clade strains, and TH1 (See Fig. 8 in the main text). However, the ID10 1aii, ID10 1bi, and region 2 have numerous differences from consensus in the second repeat and were not considered likely to encode CRISPR RNA.

### Protospacers and self-targeting immunity

Four of the spacers from the *Xenorhabdus* strains tested have self-targeting protospacers: Kenyan clade spacer 1b-1 has a self-targeting protospacer in a *palA/fhaB* gene (e.g., Kalro JASDYB01_14372); Kenyan clade spacer 1b-3 has a self-targeting protospacer in an ABC transporter-related protein (e.g., Kalro JASDYB01_13589); TH1 spacer 1a-5 has a self-targeting protospacer in *folD* (XTH1_v2_2708); and BMMCB spacer 1a-3 has self-targeting protospacers in homologs present in two predicted phage regions (LDNM01_v1_420008 and LDNM01_v1_430003).

We used a “guilt by association” approach to identify potential self-targeting immunity genes [2]. Using the MaGe comparative genome platform, we searched the genomic regions nearby protospacer genes for small ORFs that are present in self-targeting genomes but are absent in the other analyzed genomes.

*palA/fhaB*

In the Kenyan clade genomes, but not in ID10, HGB2511, or BMMCB, we found a cluster of four small ORFs encoded adjacent to the protospacer-containing *palA/fhaB* gene (See Fig. 8c in main text). These are predicted to encode a predicted *fhaB* fragment (JASDYB01_14368), a HTH cro/C1-type domain-containing protein (JASDYB01_14369), a *symE* toxin homolog (JASDYB01_14370), and a DUF2247 domain-containing protein (JASDYB01_14371). The DUF2247 gene fulfills the criteria to be an Acr candidate, since it is less than 200 aa (171aa), is encoded in the same orientation and downstream of the protospacer-encoding gene and is within four ORFs of an HTH domain-containing gene (JASDYB01_14369) that is predicted to function as an “Aca” transcriptional regulator [2]. DUF2247 is also known as “imm38” and is found in poly-immunity loci [3].

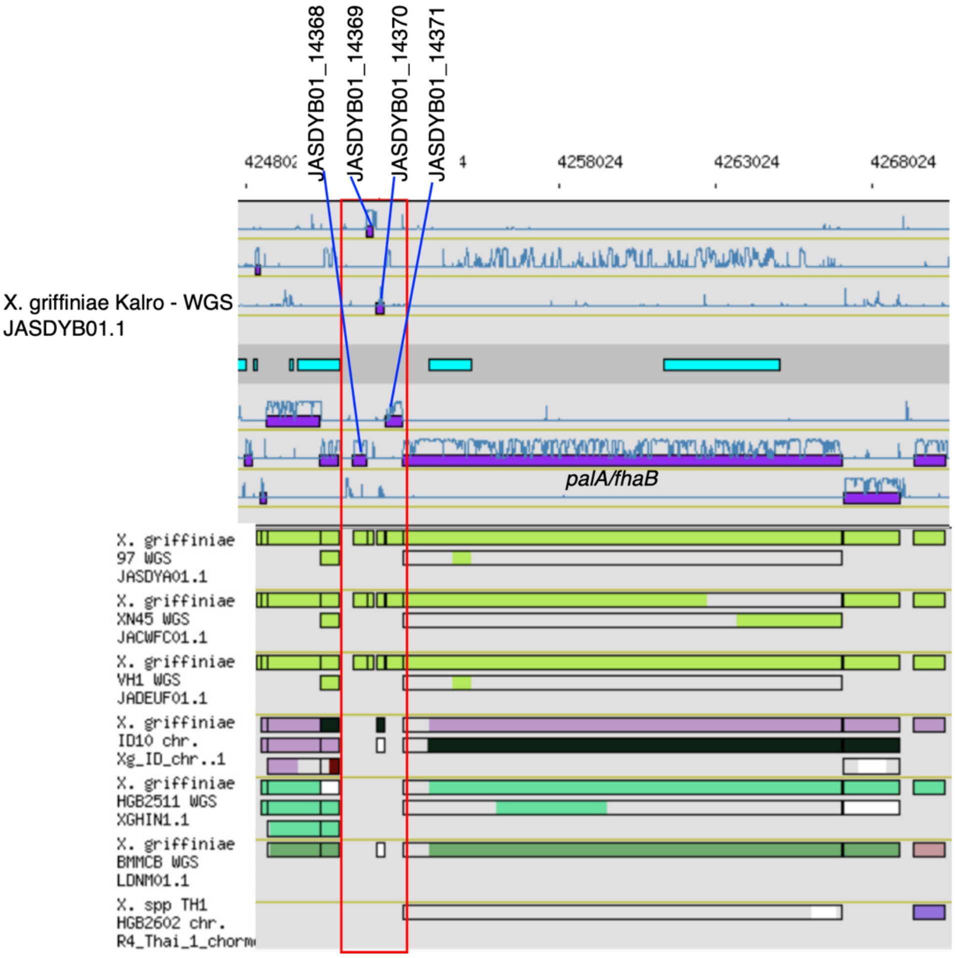


*ABC transporter-related protein*

For the Kenyan subclade spacer 1b-3, no clear evidence for Aca/Acr genes was observed in the genomic region surrounding the protospacer-containing genes (e.g., JASDYB01_13589), predicted to encode ABC transporter-related proteins. Small open reading frames (JASDYB01_13598: 65 Kda; JASDYB01_13597: 113aa, cupin domain containing; JASDYB01_13596: 51 aa; JASDYB01_13595: 39 aa) were present 3-5 Kb away from the protospacer-containing gene, but none was predicted to include an HTH-domain indicative of an Aca.

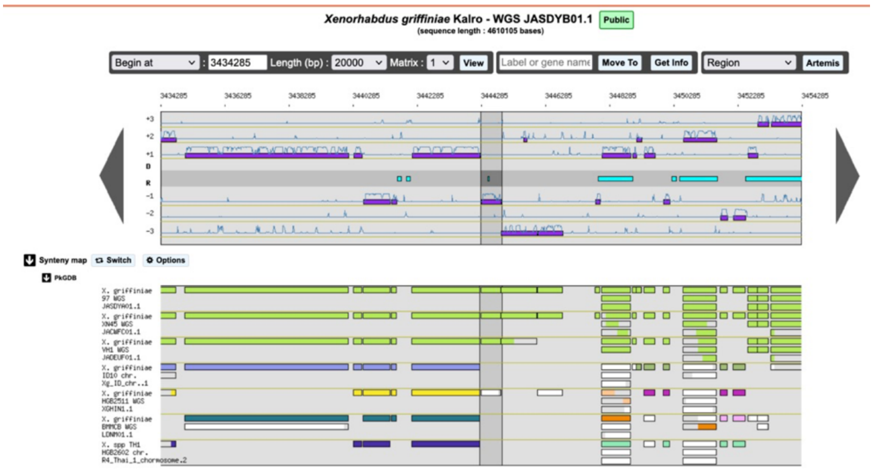


*folD*

In the TH1 genome, but not the other analyzed genomes, we found three small ORFs encoded in the vicinity of the *folD* protospacer-containing gene (bright green block arrow) with identity to TH1 spacer 1a-5. One of these, XTH1_v2_2706, is predicted to encode an integrase (black block arrow), while the other two, XTH1_v2_2705 and _2707 (pink block arrows) are predicted to encode proteins of unknown function of 38 aa and 71 aa, respectively. Neither is annotated as having an HTH-domain. Both HGB2511 and TH1 *folD* were flanked by *arg* tRNA and a S22 rRNA loci, but upstream of the *arg* tRNA they diverged. HGB2511 had phage-related genes (orange block arrows) and TH1 had a large locus (beginning of which is shown as blue lined block arrow) predicted to encode a non-ribosomal peptide synthetase.

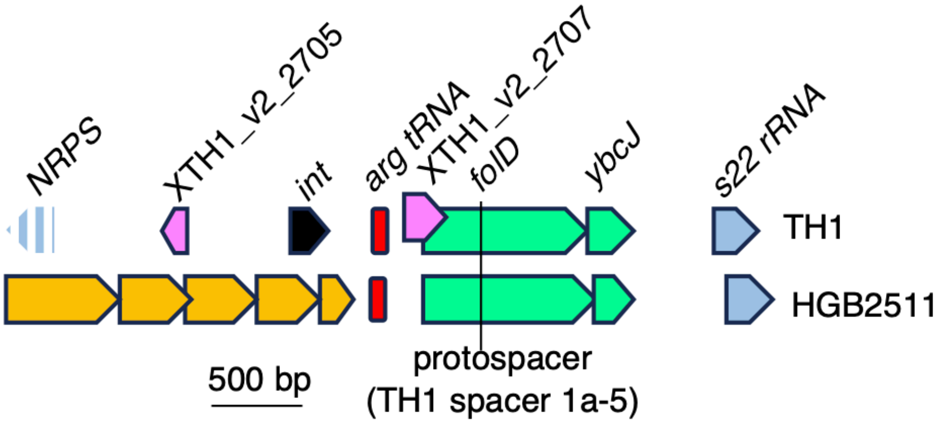

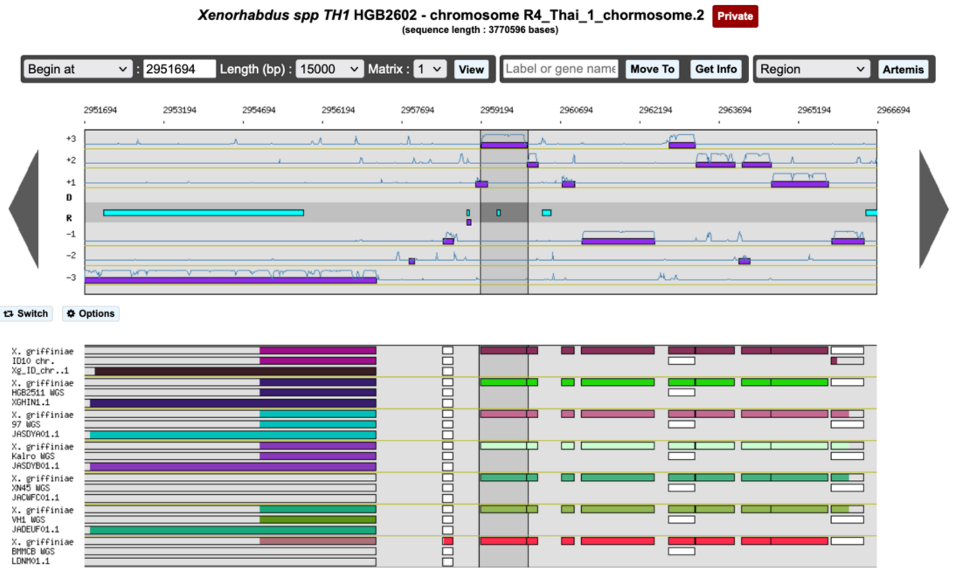
